# Sugar metabolism drives shoot branching by repressing *BRC1* through *miR319*-targeted TCP4 in *Rosa*

**DOI:** 10.64898/2026.09.09.750302

**Authors:** Léo Gouaille, Julie Mallet, Julie Legrix, Alexis Porcher, Laurent Ogé, Thibaut Perez, Johann Kraft, Maria-Dolores Pérez-Garcia, Nathalie Leduc, François Barbier, Patricks Laufs, Soulaiman Sakr, José Le Gourrierec

## Abstract

The coupling between sugar availability and shoot branching is well established, yet the underlying molecular mechanisms remain largely unknown. Here, using genome-wide profiling, we characterized the sugar signalling pathways by identifying 10 miRNAs repressed by glycolysis, the tricarboxylic acid (TCA) cycle, and/or the oxidative pentose phosphate pathway (OPPP) in rose buds. Focusing on *RhmiR319*, a miRNA repressed under active sugar metabolism, we demonstrate that disruption of both glycolysis/the TCA cycle and the OPPP induces the *RhmiR319* accumulation, leading to the concomitant repression of its targets, *RhTCP4a* and *RhTCP4b*. We additionally reveal that the *miR319/TCP4* module regulates shoot branching in *Arabidopsis*, as *miR319* overexpression impairs shoot branching, whereas TCP4 promotes it. Altered branching patterns in Rose and *Arabidopsis* were associated with the expression of the branching integrator *BRC1*. Finally, we provide molecular evidence supporting that RhTCP4a/b directly binds to the *RhBRC1* promoter and represses its transcription. Together, these results establish the *miR319– TCP4–BRC1* regulatory module as a key component in the regulation of shoot branching in response to sugar status. We propose that the *miR319-TCP4-BRC1* module is conserved between annual and perennial species and may integrate multiple hormonal and metabolic signals to shape shoot branching in plants.

## Introduction

Shoot branching is a major determinant of plant architecture (Hallé and Oldeman, 1970) that strongly influences plant fitness, crop productivity and the quality of ornamental plants (Boumaza *et al*., 2010; Xing and Zhang, 2010; Mathan *et al*., 2016). The formation of new branches depends on the release of axillary buds from dormancy (Thimann and Skoog, 1933), which is controlled by the integration of intricate hormonal, environmental and metabolic cues (Domagalska and Leyser, 2011; Liu *et al*., 2024). Branching is regulated by apical dominance, a process by which the apex inhibits the outgrowth of lower axillary branches. Apical dominance involves auxin produced by the shoot apex, which acts as an indirect regulator, as shoot-derived auxin does not directly enter axillary buds (Gälweiler *et al*., 1998; Ljung *et al*., 2002). Instead, downstream auxin-mediated signals, particularly cytokinins (CKs) and strigolactones (SLs), function antagonistically to promote or inhibit bud outgrowth, respectively (Dun *et al*., 2012; Lopez-Obando *et al*., 2015; Rameau *et al*., 2015; Beveridge *et al*., 2023).

Sugars produced by photosynthesis provide the primary carbon and energy supply sustaining plant growth and development, but they also play a decisive role in controlling shoot branching (Sakr *et al*., 2018; Doidy *et al*., 2024; Wang *et al*., 2025*c*). For instance, increased sugar availability promotes shoot branching as observed under elevated CO_2_ in *Arabidopsis* (Otori *et al*., 2017) and in genotypes with enhanced local sugar accumulation (Liu *et al*., 2019*b*; Li *et al*., 2024). Additionally, physiological studies highlight the tight coupling between sugar availability and bud outgrowth. In sorghum, reduced photosynthetic leaf area significantly inhibits bud growth (Kebrom and Mullet, 2015), whereas in wheat *tiller inhibition (tin)* mutants sugar redirection towards elongating internodes deprives axillary buds and prevents their activation (Kebrom *et al*., 2012). In pea, (Mason *et al*., 2014) demonstrated that apical dominance largely relies on carbon competition between the growing shoot apex and axillary buds, with the former acting as a dominant sink restricting sugar availability to developing axillary buds. Isotope-labelling and sugar quantification revealed that carbon and sugar content rise rapidly after decapitation, before any detectable auxin depletion in the adjacent stem. These findings support the view that sugars act as early signals triggering the release of axillary buds from apical dominance in pea (Mason *et al*., 2014; Fichtner *et al*., 2017). Consistently, the requirement of sugars in bud outgrowth has been demonstrated in one-node cutting systems (Henry *et al*., 2011; Barbier *et al*., 2015; Fichtner *et al*., 2017; Cao *et al*., 2023), in agreement with the conserved signatures of sugar starvation observed in dormant buds in multiple species (Martín-Fontecha *et al*., 2018).

Beyond their metabolic role, sugars also act as signalling molecules that control axillary bud outgrowth (Doidy *et al*., 2024; Wang *et al*., 2025*c*). Trehalose-6-phosphate (Tre6P), a key proxy of sugar status (Göbel and Fichtner, 2023), accumulates rapidly in axillary buds after decapitation and triggers shoot branching (Fichtner *et al*., 2017, 2021*a*). In parallel, disturbing the glucose-sensing pathway mediated by HEXOKINASE1 (HXK1) mutation alters shoot branching in *Arabidopsis thaliana*, showing that HXK1-dependent sugar signalling links cellular sugar status to shoot branching regulation (Barbier *et al*., 2021). In rose, the activation of the glycolysis/tricarboxylic acid cycle (glycolysis/TCA cycle) and the oxidative pentose phosphate pathway (OPPP), tightly correlates with the acquisition of bud outgrowth competence (Wang *et al*., 2021*a*). These findings position sugars as key metabolic and signalling regulators of axillary bud outgrowth and shoot branching.

At the regulatory level, *BRANCHED1* (*BRC1*)/*TEOSINTE BRANCHED1* (*TB1*), a TCP family transcription factor, acts as a master repressor of shoot branching, by integrating nutritional and hormonal cues (Aguilar-Martínez *et al*., 2007; Wang *et al*., 2019*a*; Van Es *et al*., 2024), although Tre6P promotion of bud outgrowth is independent of *BRC1* (Fichtner *et al*., 2024). Elevated sugar availability represses *BRC1*/*TB1* expression (Mason *et al*., 2014; Barbier *et al*., 2015; González-Grandío *et al*., 2017) and counteracts auxin-mediated *BRC1* induction and thereby enables bud activation across species (Wang *et al*., 2021*a*).

The molecular mechanisms linking nutritional cues to *BRC1* regulation start to be uncovered. For instance, glycolysis/TCA cycle and OPPP affect *BRC1* expression through specific regulatory elements and posttranscriptional mechanisms (Wang *et al*., 2019*a*, 2021*a*). At the transcriptional level, the light- and sucrose-responsive transcription factor *ELONGATED HYPOCOTYL 5* (*HY5*) negatively regulates *BRC1* expression in tomato (Dong *et al*., 2023). Epigenetic mechanisms also contribute, as the lncRNA APOLO negatively regulates *BRC1* transcription by modulating chromatin conformation in response to shade signal (Mammarella *et al*., 2023). Post-transcriptionally, *BRC1* expression is further refined by RNA-binding proteins and small RNAs. In rose, RhPUF4, a Pumilio and FBF protein, likely negatively regulates *RhBRC1* through its 3′UTR in connection with metabolic signals from the OPPP (Wang *et al*., 2019*b*). In parallel, miRNAs, small endogenous non-coding RNAs, are known to regulate gene expression in plants and control various developmental processes, including shoot branching (Mallet *et al*., 2022; Zhao *et al*., 2025). Among them, *miR156* functions as a key integrator of carbon availability and branching regulation, as it is regulated by environmental and metabolic signals, including shade, elevated CO₂, sucrose, and T6P levels (May *et al*., 2013; Wahl *et al*., 2013; Yang *et al*., 2013; Wang and Wang, 2015; Xie *et al*., 2017). Increased *miR156* or *miR529* promotes branching by repressing SPLs (SQUAMOSA promoter binding protein-like) transcription factors, activators of *BRC1*/*TB1* expression (Jiao *et al*., 2010; Fu *et al*., 2012; Wang *et al*., 2015; Morea *et al*., 2016; Liu *et al*., 2017; Sun *et al*., 2019; Barrera-Rojas *et al*., 2023; Li *et al*., 2023; Wei *et al*., 2024). While the involvement of *miR156* in linking sugar signalling with branching control is well established, other miRNAs regulating branching, but with no reported direct link with nutritional cues, have been described. For instance, *miR393* enhances rice tillering by targeting the auxin receptors *TRANSPORT INHIBITOR RESPONSE 1* (*OsTIR1*) and *AUXIN SIGNALING F-BOX 2* (*OsAFB2*) (Li *et al*., 2016), while a *miR160*-resistant form of *AUXIN RESPONSE FACTOR 18* (*OsARF18*) reduces tiller number (Huang *et al*., 2016). In peach, *miR6288b-3p* targets *PpTCP4* (an ortholog of *BRANCHED2*), promoting brassinosteroid biosynthesis and thereby inhibiting branching (Wang *et al*., 2024). Conversely, some miRNAs act as negative regulators of branching. Loss of *miR172* function increases branch number (Lian *et al*., 2021), while *miR171* restricts bud outgrowth by repressing *GRAS* family genes across several species (Wang *et al*., 2010; Curaba *et al*., 2013; Kravchik *et al*., 2019). In rice, *miR444* limits tillering by modulating strigolactone signalling through *MADS AFFECTING FLOWERING 57* (*OsMADS57*) and *DWARF14* (*D14*) (Guo *et al*., 2013). Also, *miR319* effects appear species-dependent: it reduces tillering in rice by targeting *Gibberellin- And Abscisic Acid-Regulated MYB* (*OsGAMYB*) and *TEOSINTE BRANCHED*/*CYCLOIDEA*/*PROLIFERATING CELL FACTOR 21* (*OsTCP21*), but promotes it in wheat via regulation of *TaGAMYB3* (Zhou *et al*., 2013; Wang *et al*., 2021*b*; Jian *et al*., 2022).

By contrast, *miR319* also regulates a set of conserved targets, a subset of the *TCP*, to control multiple biological processes, such as leaf morphogenesis, secondary cell wall formation, trichome development, and responses to biotic and abiotic stresses (Palatnik *et al*., 2003; Zhang *et al*., 2006; Ori *et al*., 2007; Sun *et al*., 2017; Vadde *et al*., 2018; Cao *et al*., 2020; Fan *et al*., 2020; Fang *et al*., 2021). In *Arabidopsis*, *miR319* targets five TCP genes (*TCP2*, *TCP3*, *TCP4*, *TCP10* and *TCP24*) and promotes cell proliferation and delays cell differentiation, thereby shaping leaves (Efroni *et al*., 2008; Sarvepalli and Nath, 2011; Schommer *et al*., 2014; Challa *et al*., 2016, 2019; Koyama *et al*., 2017, 2025; Shankar *et al*., 2023). Beyond leaf development, the *miR319/TCP* regulatory module may influence shoot branching and bud activity in woody species. In *Camellia sinensis*, the release of axillary buds from dormancy is associated with decreased *miR319* expression, which permits the accumulation of its target *TCP2* (Liu *et al*., 2019*a*). Furthermore, the *TCP4* gene has recently been shown to act downstream of SLs to regulate branching by contributing to *BRC1* repression (Huang *et al*., 2026). Collectively, these findings support a conserved mechanistic framework in which *miR319* fine-tunes bud outgrowth activity and plant architecture by modulating TCP activity.

Although miRNAs are known to contribute to glucose-dependent regulation of different aspects of plant development (Duarte *et al*., 2013; Yang *et al*., 2013; Wang *et al*., 2020*b*), whether miRNAs specifically act downstream of sugar metabolism and signalling in the control of branching, and through which molecular pathways remains unknown. In this study, we first explore how sugar metabolism and signalling reshape the rose miRNAome, and second, we specifically analyse the pivotal role of *miR319* in shoot branching. Indeed, we show that *RhmiR319* levels are influenced by a range of conditions affecting sugar metabolism, and that this miRNA in turn represses the expression of two *TCP* transcription factors, *RhTCP4a* and *RhTCP4b*, which directly bind to and downregulate *RhBRC1*. Functional evidence in *Arabidopsis* demonstrates that this *miR319*-*TCP4*-*BRC1* module controls plant architecture. Together, our results reveal a novel regulatory pathway linking sugar metabolism to the transcriptional and post-transcriptional regulation of axillary bud outgrowth.

## Materials and methods

### Plant materials, growth conditions, and phenotyping

Rose (*Rosa* ‘Radrazz’) cuttings were rooted and grown under greenhouse conditions as described previously (Bertheloot *et al*., 2020; Porcher *et al*., 2020; Wang *et al*., 2021*a*; Jiang *et al*., 2025), before transfer to a growth chamber (22°C, 16/8 h light/dark, 60% relative humidity (RH), 150 µE m⁻² s⁻¹) until the “Flower Bud Visible” phenological stage.

The *in-vitro* split-plate assays were carried out as described previously (Barbier *et al*., 2015; Porcher *et al*., 2021; Wang *et al*., 2021*a*). The basal medium was supplemented with sucrose (100 mM), 2-deoxyglucose (2-DOG, 5mM), 6-aminonicotinamide (6-AN, 10 mM), or auxin 1-naphthaleneacetic acid (NAA, 1 μM), depending on the experiments. Explants were incubated for 6 days under controlled conditions (22°C, 16/8 h light/dark, 60% RH, 90 µE.m⁻².s⁻¹) during which bud outgrowth was monitored and quantified using Fiji. All *Arabidopsis thaliana* lines were in the Col-0 background. Previously described lines were used: *jaw-D*, overexpressing *AtmiR319a* (Weigel *et al*., 2000; Palatnik *et al*., 2003); *tcp4-1* (SAIL_117D02) and *tcp4/10* (SAIL_117D02 x SALK_027514) (Schommer *et al*., 2008); *soj8* (Palatnik *et al*., 2007; Rodriguez *et al*., 2010); and *35S::mAtTCP4* (Koyama *et al*., 2007). Genotypes were confirmed by PCR using primers listed in (Supplementary Table S1). The lines *35S::RhmiR319, 35S::RhmTCP4a* and *35S::RhmTCP4b* were generated for this study. Seeds were sown with sphagnum peat and brown peat (75/25, v/v; Klasmann-Deilmann®), stratified at 4°C for 48 h in darkness, and grown under controlled conditions (22/20°C, day/night, 16/8 h light/dark, 60% RH, 150 µE.m⁻².s⁻¹). Primary rosette branches were counted every two days for 20 days after bolting (main inflorescence stem > 0.5 cm) to minimize developmental variation among genotypes (Barbier *et al*., 2021; Fichtner *et al*., 2021*b*).

For transient expression assays, 5 weeks old *N. benthamiana* plants grown under greenhouse conditions (22/18°C, day/night, 16/8 h light/dark, 60% RH, 450 µE.m⁻².s⁻¹) were used.

### Plasmids construction for plant expression

Most constructs were generated using the GoldenBraid modular cloning system (Sarrion-Perdigones *et al*., 2011, 2013). *RhTCP4a* and *RhTCP4b* coding sequences (CDS) were amplified from *Rosa* genomic DNA using Phusion™ High-Fidelity DNA Polymerase (New England Biolabs®) and domesticated into pUPD2 (Supplementary Table S1). *miR319*-resistant versions (*35S::RhmTCP4a* and *35S::RhmTCP4b*) were generated by overlap extension PCR introducing silent mutations within the predicted *miR319* target site by psRNATarget (Dai *et al*., 2018). Transcriptional units were assembled in pDGB3-α1 using the CaMV 35S promoter, CDS (with or without GFP fusions), and RbcSE9 terminator. For the *BRC1* expression reporter construct, a 1973 bp fragment upstream of the *RhBRC1* start codon was cloned upstream of the *LUCIFERASE* gene into pDGB3-α1. For multigene binary constructs, the transcriptional units (pDGB3-α1) were assembled with the pDGB3-α2 (*35S::DsRed2::RbcSE9*) and assembled into the binary vector pDGB3-Ω1.

For *35S::RhmiR319a* construct, the genomic *Rhpre-miR319* sequence was amplified and cloned into the pENTR™/TOPO® entry vector (Thermo Fisher Scientific) and recombined into the pMDC32 destination vector using an LR Clonase™ reaction.

All final constructs were verified by colony PCR and sequenced before introduction into electrocompetent *Agrobacterium tumefaciens* GV3101 cells. Transformants were selected on LB medium supplemented with the appropriate antibiotics prior to plant transformation.

### Stable and transient plant transformation

For stable expression of *35S::RhmTCP4a*, *35S::RhmTCP4b*, and *35S::RhmiR319a*, *Arabidopsis* plants were transformed by floral dipping (Clough and Bent, 1998). T1, T2 and T3 positive seeds were selected either by DsRed2 fluorescence or on MS agar medium supplemented with hygromycin. For transient assays, *35S::GFP:RhTCP4a*, *35S::GFP:RhTCP4b, 35S::GFP, 35S::RhTCP4a-35S::DsRed2*, *35S::RhTCP4b-35S::DsRed2*, *35S::DsRed2*, *pRhBRC1::LUC*, were infiltrated alone or in combination into five-week-old *N. benthamiana* plants. Adjusted *Agrobacterium tumefaciens* cultures (OD₆₀₀ = 0.5) were pelleted and resuspended in infiltration buffer containing 10 mM MgCl₂, 10 mM MES (pH 5.6) and 100 µM acetosyringone. After 2 h incubation at room temperature, suspensions were infiltrated into the abaxial side of leaves using 2 mL syringes.

### Subcellular localization of RhTCP4

Subcellular localization of RhTCP4a and RhTCP4b were determined using *35S::GFP-RhTCP4a* and *35S::GFP-RhTCP4b* constructs transiently expressed in *N. benthamiana*. Infiltrated leaves (48 h after infiltration) were imaged using a confocal microscope (Eclipse Ti, Nikon) equipped with a spinning-disk module (CrestOptics X-Light V3). GFP fluorescence was detected at 509 nm following excitation with a 475 nm laser. Confocal micrographs were acquired using identical laser power and detector settings for all samples and processed using NIS-Elements AR software (Nikon).

### Luciferase transient expression assay

Luciferin signal was measured 48 h post-infiltration in 5 mm leaf discs from agroinfiltrated *Nicotiana benthamiana leaves with 35S::RhTCP4a-35S::DsRed2*, *35S::RhTCP4b-35S::DsRed2*, or *35S::DsRed2* (as negative control), together with the reporter construct *pRhBRC1::LUC*. For measurements, leaf discs were placed in 96-well plates containing 200 µL of 10 µM synthetic D-luciferin (L9504 Sigma-Aldrich®). Luminescence was recorded for 2s every 30 min for 6 hours using a FLUOStar Omega (BMG LABTECH®). A minimum of 15 discs were analysed per treatment.

### Phylogenetic Analysis

Rose TCP protein sequences were identified by similarity searches against the *Rosa chinensis* database (rosaceae.org) using *Arabidopsis thaliana* TCP2, TCP3, TCP4, TCP10 and TCP24, known *miR319* targets as queries sequences (Palatnik *et al*., 2003; Koyama *et al*., 2007) (arabidopsis.org). Multiple protein alignments were performed with ClustalW using MEGA 12.1 (Kumar *et al*., 2024). Phylogenetic relationships were inferred using the maximum likelihood method with the Jones–Taylor–Thornton model (Jones *et al*., 1992), and node support was assessed with 1000 bootstrap replicates. *Rosa* TCP proteins and their corresponding genes were named according to their closest *Arabidopsis* orthologs. *miR319* targets in *Rosa* were predicted using psRNATarget (Schema V2-2017 release; threshold = 3.5) (Dai *et al*., 2018).

### Small RNA-seq

Small RNAs enrichment was verified by Bioanalyzer. Libraries were sequenced using DNBSEQ technology (BGI) with Unique molecular identifiers (UMIs) for accurate quantification and reduced PCR duplication bias. Twenty-one-nt reads were aligned to miRBase (Kozomara *et al*., 2019) using Bowtie (Langmead *et al*., 2009). Alignments were processed with SAMtools (Li *et al*., 2009) to obtain miRNA counts. Differential expression was analysed with DESeq2 via SARTools (Varet *et al*., 2016), with a significance threshold of *P* < 0.01. In addition, small RNA reads were aligned to the *Rosa chinensis* reference genome (Hibrand Saint-Oyant *et al*., 2018) using ShortStack version 4.0 (Johnson *et al*., 2016) to identify genomic loci producing small RNAs. Raw sequencing data are available in the NCBI Sequence Read Archive under BioProject accession PRJNA1470483 (SRA runs: SRP703594).

### mRNA and miRNA expression analyses

Total RNA was extracted from frozen, ground tissues using the CTAB method (Barbier *et al*., 2019). cDNA synthesis was performed from 500 ng of total RNA for miRNAs and 1 µg for mRNAs analysis using Invitrogen™ SuperScript III kit, with oligodT (20) primers for mRNAs and stem– loop primers for miRNAs (Supplementary Table S2), as described by Varkonyi-Gasic *et al*. (2007). Diluted cDNA was subsequently used as a template for quantitative PCR (qPCR) with gene-specific primers (Supplementary Table S2) and iQ™ SYBR Green Supermix (Bio-Rad). Reactions were run in 96- or 384-well transparent plates (Bio-Rad) and analysed on a CFX Connect™ System (Bio-Rad). Relative mRNAs and miRNAs expression was determined using the 2^−ΔΔCt^ method. mRNA and miRNA expression in *Rosa* and *Arabidopsis* samples was normalized using *RhUBIQUITIN* (*RhUBC*) and *RhU6* and *AtClatrin* and *AtU6* as housekeeping reference genes, respectively (Supplementary Table S2).

### Production of recombinant RhTCP4s proteins

*RhTCP4a* and *RhTCP4b* (previously cloned into pUPD2) were transferred into pPGN-K to generate N-terminal 6×His fusion constructs. The plasmids were transformed into *Escherichia coli* Rosetta (DE3). Positive clones were grown in LB medium containing kanamycin to OD₆₀₀ 0.5, induced with 0.1 mM IPTG, and incubated overnight at 20°C. Cells were harvested, resuspended in an extraction buffer (50 mM Tris-HCl, pH 8.0, 200 mM NaCl) containing EDTA-free protease inhibitor cocktail (Roche) and lysed by sonication (Vibra-Cell™ 72434, Bioblock Scientific) at 60% amplitude for 10 cycles of 10s on / 10s off on ice. Lysates were centrifuged and the supernatant was supplemented with 10 mM imidazole and incubated with Ni-NTA agarose for 1 h at 4°C and washed with a buffer containing 40 mM imidazole. His-tagged RhTCP4a/b proteins were eluted with a buffer containing 250 mM imidazole and 10% glycerol. Protein expression and purification efficiency were assessed by SDS-PAGE and Western blot using an HRP-conjugated monoclonal anti-His antibody (Bio-Rad).

### Electrophoretic mobility shift assay (EMSA)

EMSA was performed using an adapted protocol (Wang *et al*., 2022). Oligonucleotide probes containing TCP-binding motifs isolated from the *RhBRC1* promoter were synthesized with 5′DIG labels (Eurofins Genomics). Binding reactions contained 1 µM of probe and 1 µg of recombinant His-RhTCP4a or His-RhTCP4b were incubated in 100 mM HEPES, 5 mM EDTA (pH 8.0), 50 mM (NH₄)₂SO₄, 5 mM DTT, 150 mM KCl, and 1% (v/v) Tween 20, for 25 min at room temperature. Protein–DNA complexes were resolved on 6% native polyacrylamide gel at 4°C for 90 min, transferred to a HYBOND™ N+ membrane, and detected with HRP-conjugated anti-DIG antibody (Bio-Rad) with the ChemiDoc™ XRS+ System (Bio-Rad). Specificity was tested by competition assay using 10x and 50x-excess unlabelled wild-type or mutated oligonucleotides. All oligonucleotides are listed in Fid. 6d.

### Statistical analyses

All data were analysed using Student’s t-test for pairwise comparisons or Kruskal-Wallis test for multiple-group comparisons followed by Dunn’s post hoc test. A significance level of *P* < 0.05 was used. Significant differences are indicated by asterisks (*) or letters on the graphs. Data are expressed as mean ± standard deviation (SD) or standard error of the mean (SEM), as indicated in the figure legends.

## Results

### The glycolysis/TCA cycle and OPPP repress *RhmiR319-3p* accumulation

Our previous data indicate that both the glycolysis/TCA cycle and the OPPP are necessary to promote rose bud outgrowth in response to sugar availability (Wang *et al*., 2021*a*). To identify microRNAs acting downstream of sugar metabolism and signalling, we performed a small RNA-seq on rose axillary buds treated with inhibitors of glycolysis/TCA cycle (2-DOG) (Jang and Sheen, 1994; Lejay *et al*., 2003; Rabot *et al*., 2012; Xiong *et al*., 2013; Stitz *et al*., 2023), and OPPP (6-AN) (Köhler *et al*., 1970; Lejay *et al*., 2008; Wang *et al*., 2021*a*), alone or combined, under high sucrose (100 mM).

Quantifying bud outgrowth over a 6 days confirmed earlier findings: inhibition of glycolysis/TCA cycle or OPPP reduced bud length to approximately 0.25 and 0.3 cm at day 6, respectively, versus 0.8 cm with sucrose alone (Fig. 1A, B) and simultaneous inhibition of both pathways resulted in an even stronger repression (Fig. 1A) (Wang *et al*., 2021*a*). Next, total RNAs were extracted from samples incubated for 24 h under the same experimental conditions and subjected to small RNA-seq to identify microRNAs whose abundance is modulated by the inhibition of glycolysis/TCA cycle or the OPPP. The 24 h time point was selected because previous studies showed that numerous transcripts display significant changes at this stage, before bud elongation (Girault *et al*., 2010; Rabot *et al*., 2012; Barbier *et al*., 2015; Porcher *et al*., 2021; Wang *et al*., 2021*a*; Jiang *et al*., 2025).

**Fig. 1.**
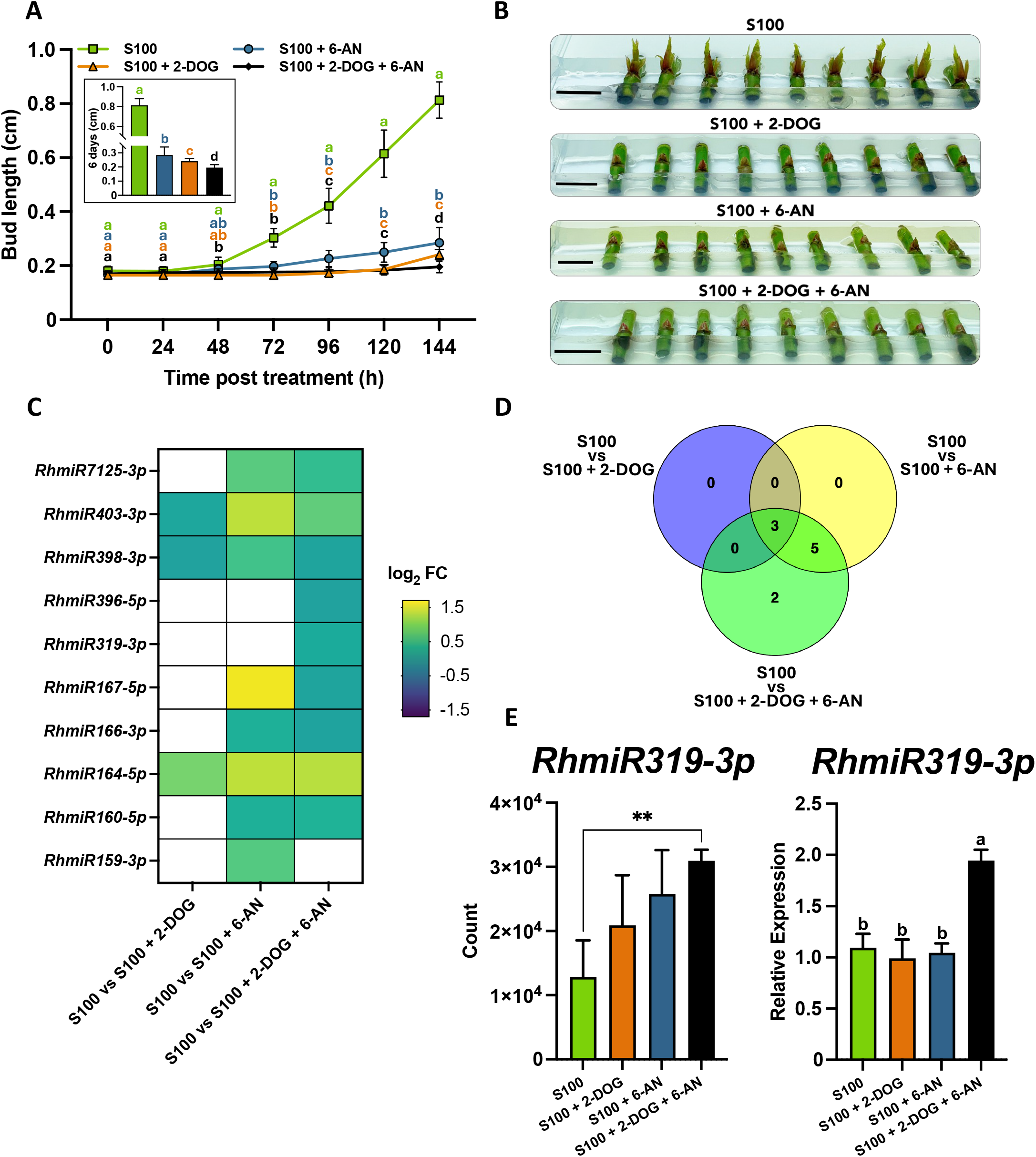
The glycolysis/TCA cycle and OPPP regulate *RhmiR319* levels in rose axillary buds. (A) Growth kinetics of *in vitro* cultured buds incubated with 100 mM sucrose (S100) and supplemented with 5 mM glycolysis/TCA cycle inhibitor (2-DOG) or 10 mM OPPP inhibitor (6-AN) for 6 days (*n* ≥ 9). (B) Representative photographs showing buds grown on different culture media after 6 days. Bars, 1 cm. (C) Heatmap showing differentially expressed miRNAs across the indicated conditions. Coloured cells represent significantly differentially expressed miRNAs (*P* < 0.01). (D) Venn diagram representing numbers of differentially expressed miRNAs across the different experimental conditions. (E) *RhmiR319* levels in axillary buds in response to the different treatments. *RhmiR319* levels represented in counts (variance-stabilized transformation, VST) based on miRNA-seq data (left panel) or obtained by qRT-PCR (right panel). Data represent the mean ± SD. In (A) and (E), the letters indicate significant differences (*P* < 0.05) based on a non-parametric Kruskal–Wallis test followed by Dunn’s post hoc test. In (E) Asterisks indicate significant differences according to Student’s *t*-test (**, *P* < 0.01).

Small RNA-seq analysis identified 10 up-regulated miRNAs (*RhmiR159-3p*, *RhmiR160-5p*, *RhmiR164-5p*, *RhmiR166-3p*, *RhmiR167-5p*, *RhmiR319-3p*, *RhmiR396-5p*, *RhmiR398-3p*, *RhmiR403-3p* and *RhmiR7125-3p*) in response to the glycolysis/TCA cycle (2-DOG) and/or the OPPP (6-AN) inhibitors (Fig. 1C). Most miRNAs -nine out of ten-were upregulated under the combined blockage of glycolysis/TCA cycle and the OPPP. Among them, two were regulated exclusively upon combined inhibition of the two pathways (*RhmiR319-3p* and *RhmiR396-5p*), three were upregulated by all treatment conditions (*RhmiR164-5p*, *RhmiR398-3p* and *RhmiR403-3p*), and four were upregulated by combined inhibition of the two pathways or by the inhibition of the OPPP pathway alone (*RhmiR160-5p*, *RhmiR167-5p*, *RhmiR7125-3p* and *RhmiR166-3p*) (Fig. 1D). Altogether, transcriptomic analyses indicated that the glycolysis/TCA cycle and the OPPP pathways modulate miRNA accumulation in rose axillary buds, mainly in a combined manner or specifically via the OPPP pathway.

Among the 10 miRNAs identified, we focused on *RhmiR319*, which was specifically up-regulated upon combined inhibition (S100 + 2-DOG + 6-AN), a result confirmed by qRT–PCR, with a 2-fold change relative to the control (S100) (Fig. 1E). This miRNA was prioritized because it is induced by darkness in rose axillary buds (Mallet, 2022), a condition tightly linked to sugar starvation in buds (Girault *et al*., 2010; Henry *et al*., 2011; Rabot *et al*., 2012, 2014).

### Expression of *RhmiR319*-targeted *RhTCP4a* and *RhTCP4b* is promoted by the glycolysis/TCA cycle, and OPPP pathways

To identify *TCP* genes targeted by *miR319* in rose, miRNA target prediction was performed using the psRNATarget tool (Dai *et al*., 2018) probing *RhmiR319* against the rose genome. In parallel, known *miR319*-targeted *Arabidopsis TCP* genes (*TCP2*, *TCP3*, *TCP4*, *TCP10* and *TCP24*) were used as queries in BLAST searches to identify their orthologs in the rose genome and phylogenetic analysis of full-length protein sequences. These analyses identified one ortholog of *AtTCP2* (*RhTCP2*), one ortholog of *AtTCP10* (*RhTCP10*), and three co-orthologs of *AtTCP4*, named *RhTCP4a*, *RhTCP4b*, and *RhTCP4c*, in rose (Fig. 2A). No orthologs of *AtTCP3* nor *AtTCP24* were detected in the rose genome (Fig. 2A). Multiple sequence alignment further showed that the TCP domain is highly conserved between *Arabidopsis* and rose TCP proteins, supporting their evolutionary and functional relationship (Supplementary Fig. S1A). Target prediction analysis further revealed that *RhTCP2*, *RhTCP4a*, and *RhTCP4b* contained a predicted *RhmiR319* binding site, whereas *RhTCP4c* and *RhTCP10* lack such predicted sites, consistent with C-terminal truncation, the region that usually contains the *miR319* binding site (Fig. 2B, S1B).

**Fig. 2.**
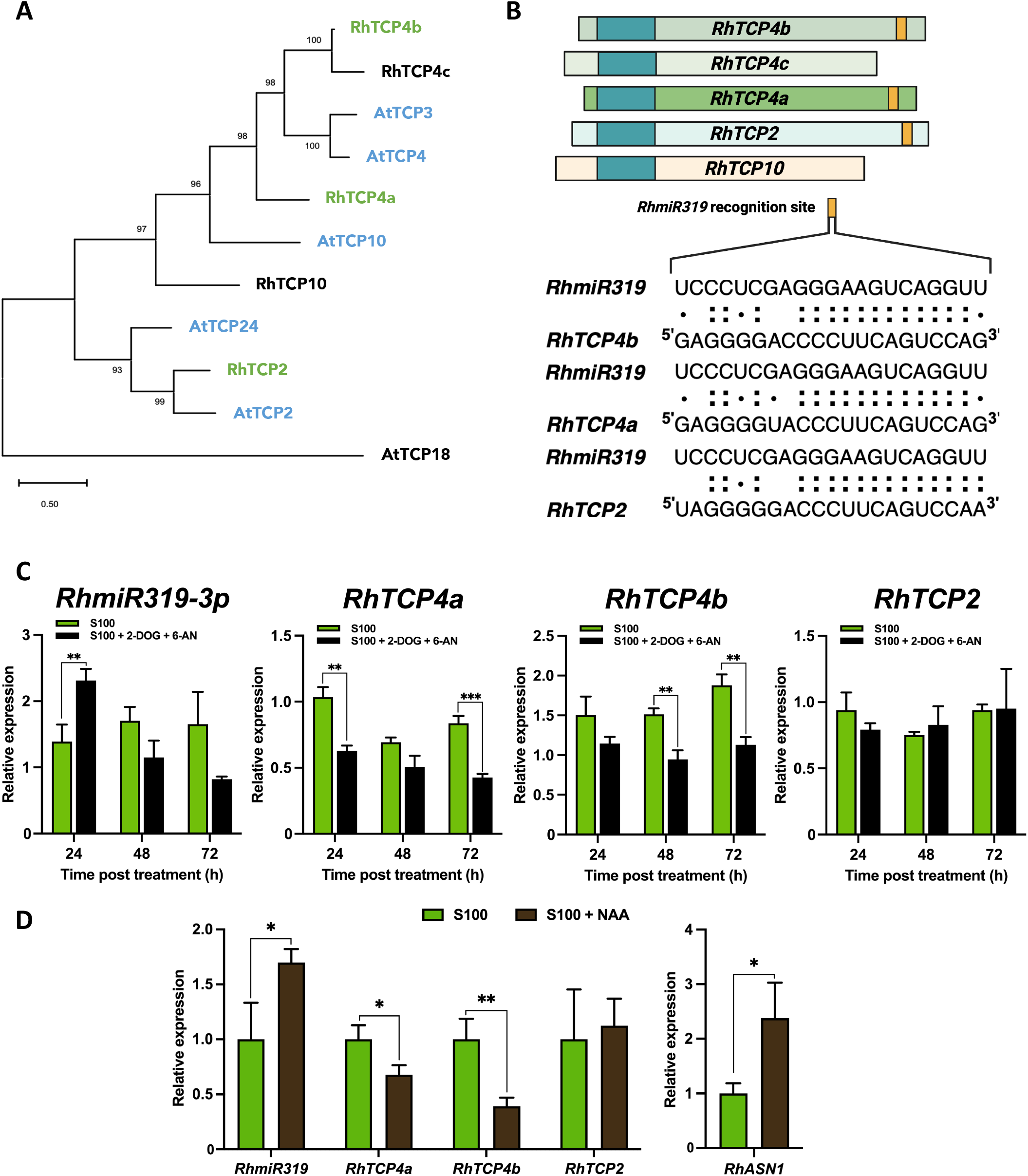
The glycolysis/TCA cycle and OPPP regulate the *RhmiR319* targets *RhTCP4a* and *RhTCP4b* at the transcriptional level in rose axillary buds. (A) Phylogenetic tree of predicted TCP targets of *miR319* in *Arabidopsis thaliana* (in blue) and their orthologs in *Rosa* (in green). Black-coloured TCPs are Rosa genes not targeted by *miR319*. The tree was rooted with AtTCP18 as the outgroup. (B) Schematic representation of *RhTCP2*, *RhTCP4a*, *RhTCP4b*, *RhTCP4c*, and *RhTCP10* transcripts, the TCP domain is indicated in blue and the *RhmiR319* binding site in orange. Below, the complementarity of sequences between *TCPs* and *RhmiR319* is represented by double points for Watson-Crick base pairing and a single point for G-U wobbles. (C) Relative expression levels of *RhmiR319*, *RhTCP4a*, *RhTCP4b*, and *RhTCP2* target genes in axillary buds incubated with 100 mM sucrose alone (S100) and supplemented with 5 mM 2-DOG and 10 mM 6-AN (S100 + 2-DOG + 6-AN). (D) Relative expression levels of *RhmiR319, RhTCP4a*, *RhTCP4b*, *RhTCP2* target genes and *RhASN1* in axillary buds incubated for 24 hours with 100 mM sucrose alone (S100) or supplemented with 1 µM auxin (S100 + NAA). Data represent the mean ± SD from three biological replicates. Asterisks indicate significant differences according to Student’s *t*-test (**, *P* < 0.01; ***, *P* < 0.001).

We then investigated how glycolysis/TCA cycle- and OPPP-mediated sugar signalling affect *RhmiR319*/*RhTCP2*, *RhTCP4a*, and *RhTCP4b* module expression during rose axillary bud growth. To address this, expression levels of this miRNA and its mRNA targets were quantified by qRT-PCR on single nodal rose segments grown in split plates in the presence of 100 mM sucrose (S100), alone or in combination with the 2-DOG and 6-AN inhibitors, over a 72 h time course (S100+2-DOG+6-AN).

Combined inhibition of glycolysis/TCA cycle and the OPPP pathway caused approximately 1.7-fold induction of *RhmiR319* at 24 h (Fig. 2C), in line with our earlier findings (Fig. 1E). Conversely, the levels of the predicted target genes, *RhTCP4a* and *RhTCP4b*, were significantly reduced at 24 and 48 hours, respectively, while the levels of *RhTCP2* were unchanged (Fig. 2C). Indirect repression of these two sugar metabolism pathways by auxin (Wang *et al*., 2021*a*) produced the same patterns: increased *RhASN1* (*ASPARAGINE SYNTHETASE1*), a sugar starvation marker (Lam *et al*., 1994; Wang *et al*., 2021*a*), elevated *RhmiR319* transcript levels, reduced expression of *RhTCP4a* and *RhTCP4b*, and unchanged expression of *RhTCP2* (Fig. 2D). These results support a link between auxin, sugar deficiency, and *miR319*-mediated bud repression. The anti-correlated expression levels between *RhmiR319* and *RhTCP4a/b* were further confirmed by a transient heterologous expression experiment in *N. benthamiana* leaves and this repression is partially alleviated when silent mutations were introduced in the miRNA recognition sites (Supplementary Fig. S2A, B).

Taken together, these results indicated that the sugar metabolism regulates *RhmiR319*/*RhTCP4a*-*RhTCP4b* modules through the glycolysis/TCA cycle pathway and the OPPP during rose bud outgrowth.

### Loss of apical dominance regulates the *RhmiR319*/*RhTCP4a*-*RhTCP4b* modules *via* sugar metabolism and signalling

To enlarge our conclusions gained from the *in vitro* study of excised buds to the whole-plant level, we assessed the expression levels of *RhmiR319* and its TCP targets in dormant buds every 3 hours over a 24-hour cycle. *RhmiR319* expression is induced early in the day, peaking 3 hours after the start of the photoperiod (ZT3) and then gradually decreasing throughout the day (Supplementary Fig. S3A). In contrast, the *RhTCP2*, *RhTCP4a*, and *RhTCP4b* targets showed the opposite pattern, with lowest expression at ZT3 (Supplementary Fig. S3A). The strongest association was observed for *RhTCP4a*, which displayed the highest correlation coefficient (*r* = -0.74) and the most significant *P*-value (*P* < 0.001) (Supplementary Fig. S3B).

We first confirmed that the loss of apical dominance induced axillary bud outgrowth, as evidenced by the enhanced growth of axillary buds following decapitation (Fig. 3A, B). We next analysed the effects of apical dominance loss on the regulation of the *RhmiR319/RhTCP4a-b* module. The release of apical dominance promotes bud outgrowth while enhancing carbon allocation and sugar metabolism in axillary buds (Snow, 1929; Mason *et al*., 2014; Wang *et al*., 2021*a*). To minimize circadian regulation, subsequent analyses were performed at ZT3, when *RhmiR319* expression is maximal and target genes are minimally expressed (Supplementary Fig. S3A). Plants were decapitated at ZT0, and axillary buds were harvested at ZT3 on three consecutive days (ZT3-D0, ZT3-D1 and ZT3-D2; 3 h, 3+24 h and 3+48 h after decapitation at ZT0, respectively) and compared with intact plants sampled at the same time points.

**Fig. 3.**
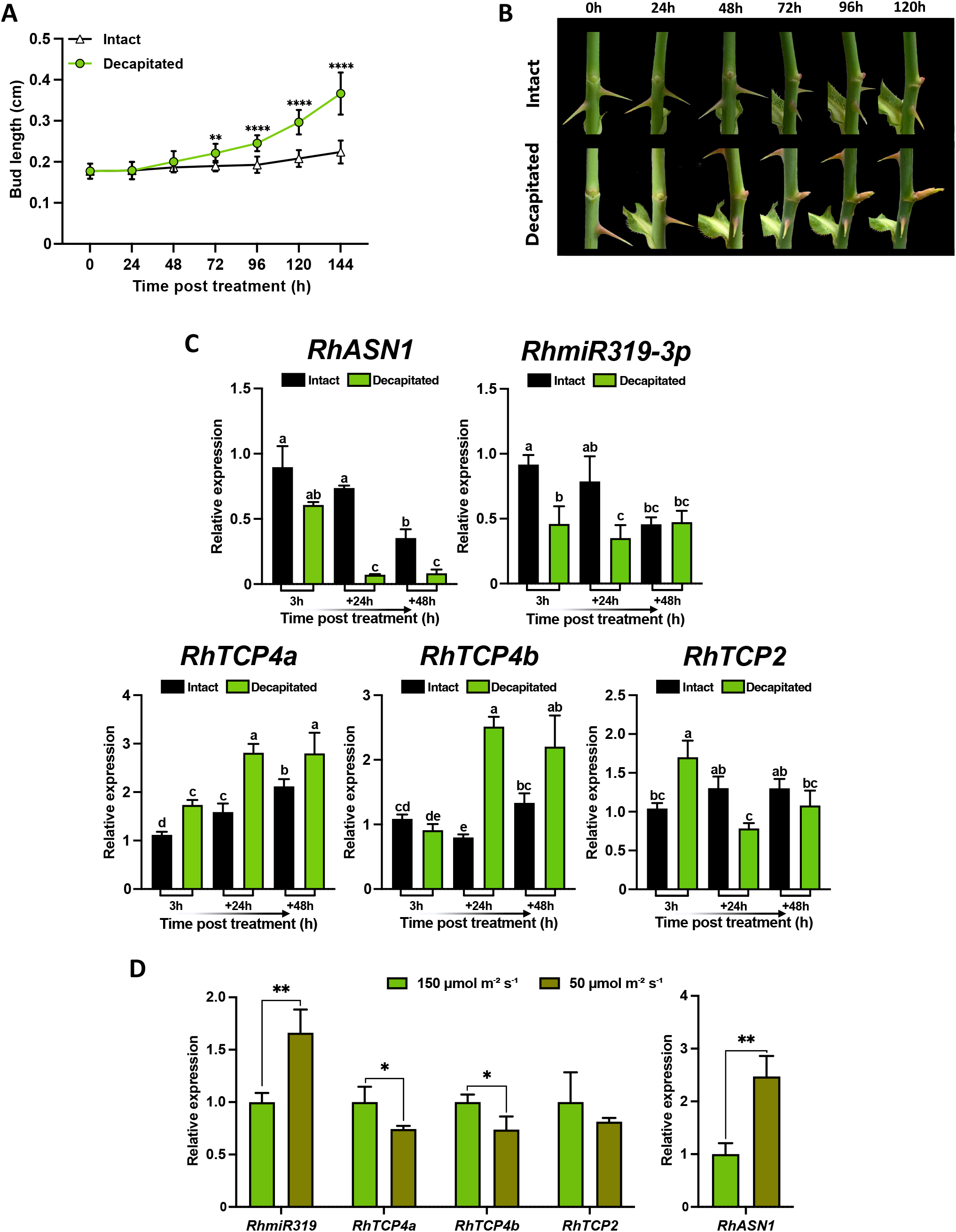
*RhmiR319*-*RhTCP4a/b* module is regulated by apical dominance under the influence of auxin. (A) Bud outgrowth kinetics in response to apical dominance release by beheading over 6 days. Data represent the mean ± SD (*n* ≥ 9). (B) Representative photographs showing buds from intact or decapitated plants after 5 days. (C) Relative expression levels of *RhASN1*, *RhmiR319*, *RhTCP4a*, *RhTCP4b*, and *RhTCP2* in axillary buds of intact and decapitated plants. (D) Relative expression levels in decapitated plants exposed to high or low light intensity (150 or 50 µmol m⁻² s⁻¹) for 6 hours. Data represent the mean ± SD from three biological replicates. In (C), letters indicate significant differences (*P* < 0.05) based on a non-parametric Kruskal– Wallis test followed by Dunn’s post hoc test. In (a,d) Asterisks indicate significant differences according to Student’s t-test (*, *P* < 0.05; **, *P* < 0.01; ****, *P* < 0.0001).

In this framework, *RhASN1* was dramatically reduced at ZT3-D1 and ZT3-D2 after decapitation (Fig. 3C), consistent with a sustained change in sugar status and carbon allocation toward axillary buds (Girault *et al*., 2010; Mason *et al*., 2014; Fichtner *et al*., 2017). Decapitation reduced *RhmiR319* expression by approximately 2-fold at ZT3-D0 and ZT3-D1 whereas no significant difference was observed at ZT3-D2 relative to intact plants. In contrast, *RhTCP4a* expression increased by 1.5- to 1.8-fold at all three time points in decapitated plants. *RhTCP4b* expression was unaffected by decapitation at ZT3-D0 but increased by approximately 3-fold at ZT3-D1 and 1.6-fold at ZT3-D2. *RhTCP2* exhibited a transient induction at ZT3-D0, followed by a significant drop at ZT3-D1, before returning to levels comparable to those of intact plants at ZT3-D2 (Fig. 3C).

To further investigate the link between sugar metabolism and the *RhmiR319*/target gene module, we modulated sugar production via photosynthesis by growing decapitated plants for 6 hours under two light intensities (150 or 50 µmol m⁻² s⁻¹). Buds from plants exposed to low light intensity (50 µmol m⁻² s⁻¹) exhibited high expression of *RhASN1*, indicative of a relative sugar deficiency state. Under these conditions, *RhmiR319* levels increased by approximately 1.7-fold, whereas *RhTCP4a* and *RhTCP4b* expression decreased to 0.74-fold. *RhTCP2* levels remained unchanged (Fig. 3D). Overall, these findings indicate that the *RhmiR319*/*RhTCP4a*-*RhTCP4b* regulatory module responds to changes in sugar availability, with *RhmiR319* acting to repress *RhTCP4a* and *RhTCP4b* in sugar-deficient conditions, thus contributing to bud outgrowth inhibition.

### *miR319* negatively regulates shoot branching in *Arabidopsis thaliana*

Our expression data in rose buds suggests that the *RhmiR319*/*RhTCP4a*-*RhTCP4b* module could regulate shoot branching. Because stable genetic manipulation of rose *in planta* is difficult, we used *Arabidopsis* as a model system to test whether *miR319* negatively regulates shoot branching. Thus, we overexpressed the rose *miR319* gene in *Arabidopsis* under the 35S promoter (*35S::RhmiR319*) and compared the obtained line to the *jaw-D* mutant which overexpressed endogenous *Arabidopsis miR319a* (Palatnik *et al*., 2003, 2007). As in *jaw-D*, the *35S::RhmiR319* line showed crinkled leaves, though the phenotype is less pronounced (Supplementary Fig. S4). Both lines also showed delayed axillary rosette leaf bud emergence, reaching their highest primary rosette branches number at approximately 12 days after bolting, whereas this stage is reached after 8 days in wild type (WT; Col-0). In addition, both lines showed an average reduction of approximately 1.5 in the final branch number compared to WT (Fig. 4A, B).

**Fig. 4.**
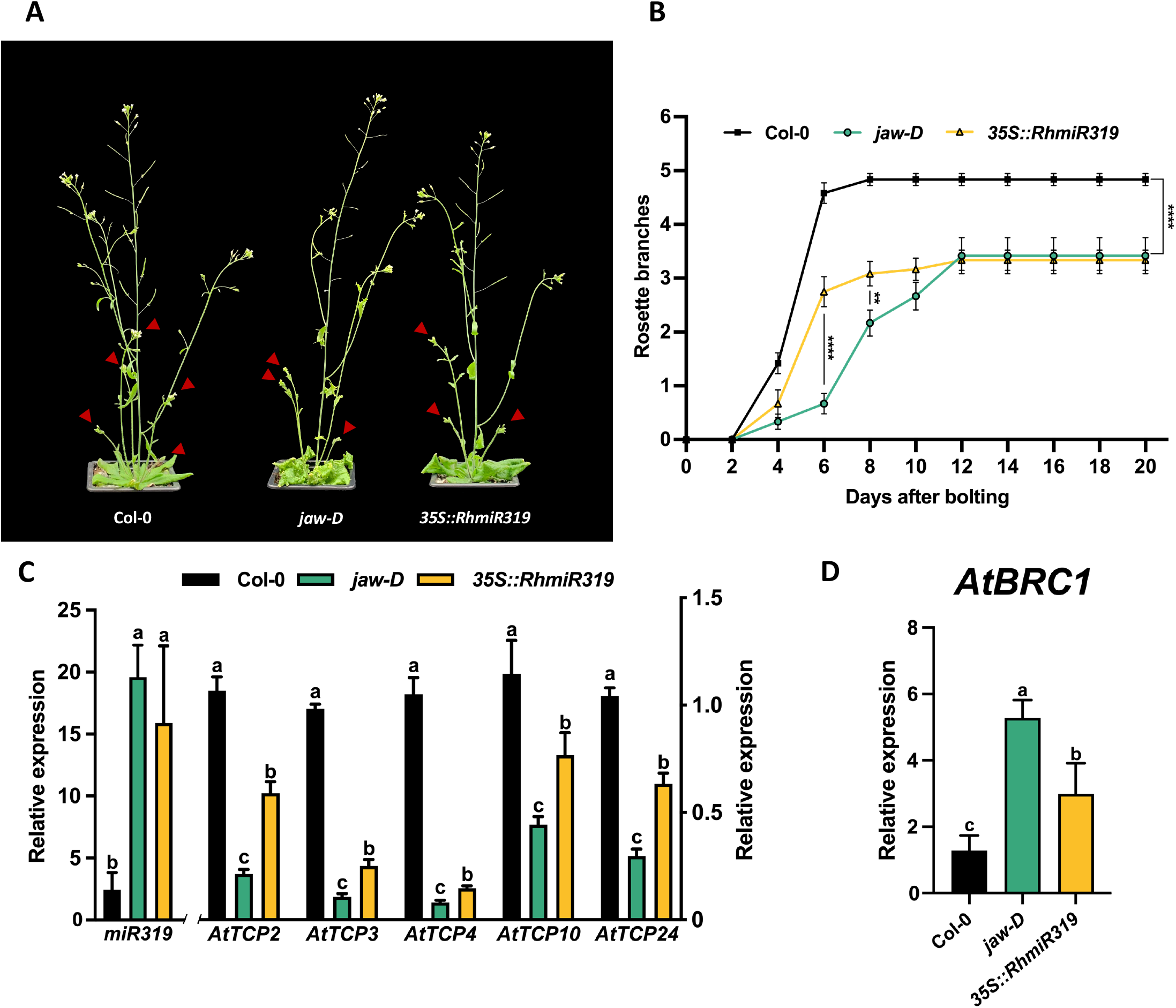
*miR319* inhibits shoot branching in *Arabidopsis*. (A) Photographs representing the branching phenotype of WT, *jaw-D* and *35S::RhmiR319* lines at 14 days after bolting. (B) Total number of primary rosette branches in the WT, *jaw-D*, and *35S::RhmiR319* lines. (C) Relative expression levels of *miR319* and its targets (*AtTCP2*, *AtTCP3*, *AtTCP4*, *AtTCP10* and *AtTCP24*) in WT, *jaw-D*, *35S::RhmiR319* lines. (D) Levels of *AtBRC1* in WT, *jaw-D* and *35S::RhmiR319* lines. In (B) data represent the mean ± SEM (*n* = 12). Asterisks indicate significant differences according to Student’s t-test (**, *P* < 0.01; ****, *P* < 0.0001). In (C,D) data represent the mean ± SD from three biological replicates. Letters indicate significant differences (*P* < 0.05) based on a non-parametric Kruskal–Wallis test followed by Dunn’s post hoc test.

To investigate whether *RhmiR319* negatively regulates the expression of the *Arabidopsis* endogenous TCP genes, qRT-PCR analysis of *miR319* and *miR319*-targeted TCP transcripts was carried out in *Arabidopsis* lines overexpressing *RhmiR319* or *jaw-D*. The overexpression of *miR319* from either the *Arabidopsis* or rose sequences led to reduced mRNA levels of the five *miR319*-targeted *TCP* genes in *Arabidopsis* (Fig. 4C), highlighting the functional conservation of *miR319* between these two species. However, *TCP* genes appeared to be more strongly downregulated in the *jaw-D* line, which correlated with the delayed lateral branch emergence and the more pronounced crinkled leaf phenotypes (Fig. 4B).

Because *BRC1* integrates hormonal and sugar inputs to control bud outgrowth (Aguilar-Martínez *et al*., 2007; Wang *et al*., 2019*a*), we sought to determine whether *miR319* could influence branching through modulation of *BRC1* expression. *AtBRC1* transcript levels were significantly higher in *jaw-D* and *35S::RhmiR319* lines compared with the WT (Fig. 4D). These results indicated that reduced branching in *miR319*-overexpressing lines correlates with an increase of *AtBRC1* expression.

### TCP4 increases shoot branching in *Arabidopsis*

Since TCP4 appears to be involved in branching regulation in rose, we examined whether changes in *TCP4* expression could account for the branching phenotypes observed in *Arabidopsis*. First, we focused on the role of AtTCP4 in *Arabidopsis,* as RhTCP4a/b are involved in rose bud outgrowth (Fig. 2C, 3C) and their expression was one of the most strongly reduced in both *miR319*-overexpressing lines (Fig. 4C). To do this, we scored primary rosette branches in single *tcp4,* as well as *tcp10* and double *tcp4*/*10* loss-of-function mutants as a functional redundancy was previously reported for these two genes during petal greening (Zheng *et al*., 2022). While single *tcp4* or *tcp10* mutants exhibited a number of branches similar to that of the WT, the *tcp4/10* double mutant showed a significant reduction in lateral branch number, with approximately one fewer branch per plant compared to the WT (Fig. 5A, B and supplementary Fig. S5). To determine whether the branching phenotype of *tcp* mutants was associated with altered *AtBRC1* expression, we measured *AtBRC1* expression in WT and in *tcp4* and *tcp4/10* mutants. *AtBRC1* expression remained unchanged in the *tcp4* mutant but was significantly increased in the *tcp4*/*10* double mutant (Fig. 5C). These results indicate that reduced branching in *tcp4/10* double mutants correlates with increased *AtBRC1* expression.

**Fig. 5.**
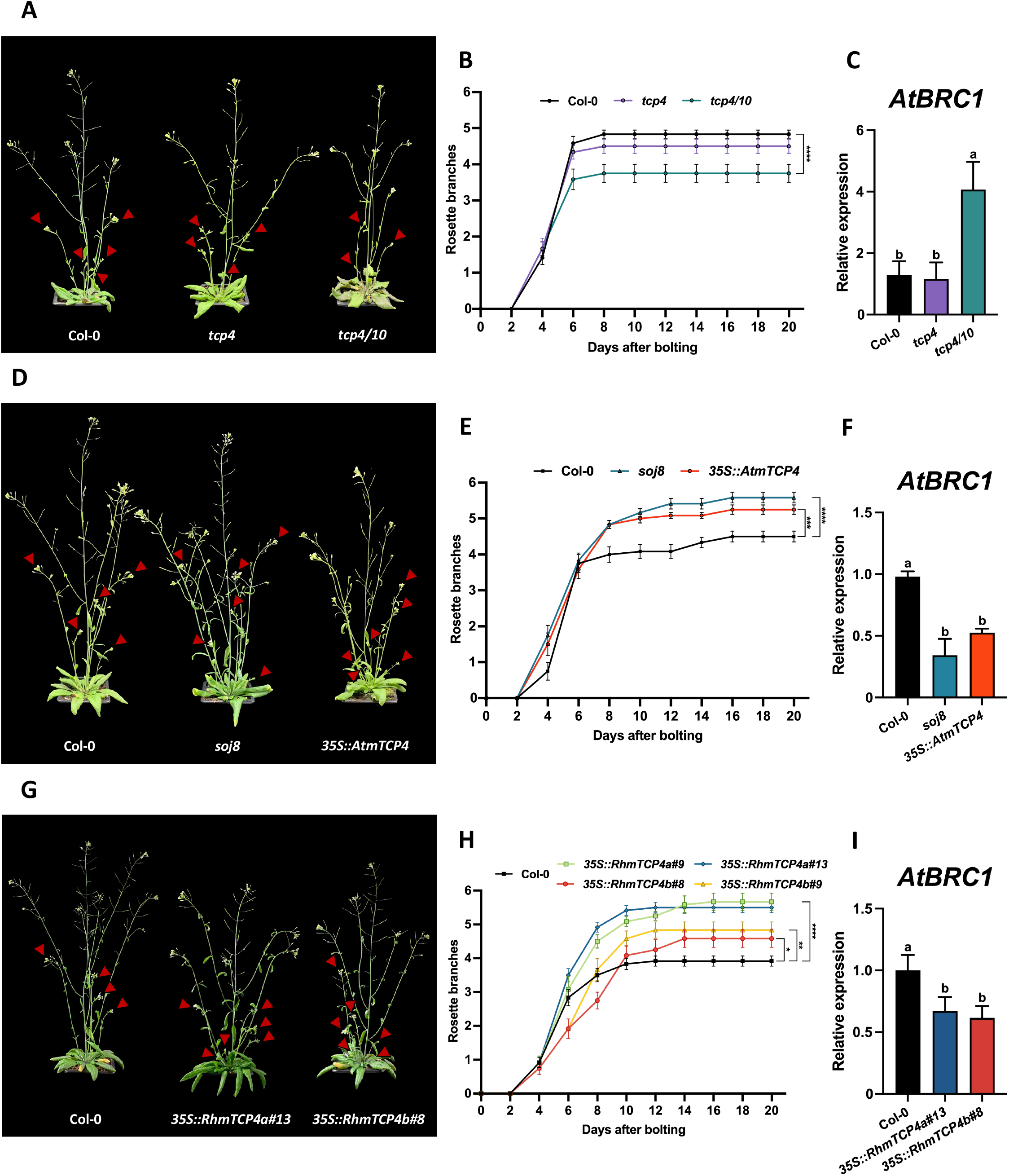
TCP4 promotes shoot branching in *Arabidopsis*. (A,D,G) Photographs representing the branching phenotype of WT, *tcp4*, *tcp4/10, soj8*, *35S::AtmTCP4* and *35S::RhmTCP4a/b* lines at 14-days after bolting. Total number of primary rosette branches in (B) the WT, *tcp4*, *tcp4/10*, (E) *soj8 and 35S::AtmTCP4* and (H) *35S::RhmTCP4a/b* lines. Levels of *AtBRC1* in (C) WT, *tcp4* and tcp4/10 lines, (F) WT, *soj8*, and *35S::AtmTCP4* lines and (I) WT and *35S::RhmTCP4a/b* lines. In (B,E,H), data represent the mean ± SEM (*n* = 12). Asterisks indicate significant differences according to Student’s *t*-test (***, *P* < 0.001; ****, *P* < 0.0001). In (C,F,I) data represent the mean ± SD from three biological replicates. Letters indicate significant differences (*P* < 0.05) based on a non-parametric Kruskal–Wallis test followed by Dunn’s post hoc test.

Next, to determine whether increasing *TCP4* levels is sufficient to stimulate branching in *Arabidopsis*, we analysed two mutant lines. The *soj8* mutant which carries a silent point mutation in the *miR319* target site of *AtTCP4*, reducing *miR319* binding (Palatnik *et al*., 2007), and *35S::AtmTCP4*, which constitutively expresses a *miR319*-resistant *TCP4* gene (Koyama *et al*., 2007). In contrast to the *tcp4/10* loss-of-function mutant, both *miR31*9-resistant *TCP4* lines showed a significant increase in primary rosette branching compared with WT plants, with approximately one additional branch per plant (Fig. 5D, E). *AtBRC1* expression was reduced in both *soj8* and *35S::AtmTCP4* lines compared with WT, indicating that increased TCP4 activity is associated with decreased *AtBRC1* expression (Fig. 5F). In parallel, to test the effect of the rose *miR319* targets *RhTCP4a*/*b*, we overexpressed *miR319*-resistant versions of rose *RhTCP4s* (*35S::RhmTCP4a* and *35S::RhmTCP4b*) in *Arabidopsis*. These constructs led to a significant increase in the final number of primary rosette branches compared with WT, with an average increase of approximately 1 branch for *RhmTCP4b* lines and 1.5 branches for *RhmTCP4a* lines (Fig. 5G, H and supplementary Fig. S6). Consistent with *soj8* and *35S::AtmTCP4* lines, *AtBRC1* expression was also significantly reduced in both *35S::RhmTCP4a* and *35S::RhmTCP4b* lines, further supporting a negative relationship between TCP4 activity and *AtBRC1* expression (Fig. 5I and supplementary Fig. S6A, B).

Altogether, our loss- and gain-of-function analyses support a positive role of TCP4 in the regulation of shoot branching in *Arabidopsis*. Consistent with the enhancement branching phenotype previously reported in triple *tcp* mutants (*tcp3*/*4*/*10*) (Huang *et al*., 2026), *AtBRC1* expression increased in *tcp4*/*10* backgrounds. Conversely, *AtBRC1* expression was consistently downregulated in both *35S::AtmTCP4* and *soj8* backgrounds. Although this contrasts with observation from a previously analysed *mTCP4* line (Huang *et al*., 2026), the concordant molecular and phenotypic responses observed in independent TCP mutants (Fig. 5D, E), supports a model in which TCP4 promotes shoot branching, at least partly through a modulation of *BRC1* expression. These findings further suggest that the TCP-dependent shoot branching regulation may depend on the genetic and developmental context.

### RhTCP4a/b repress transcription of *RhBRC1* by binding to its promoter

Previous work has shown that *AtTCP4* can directly bind the *AtBRC1* promoter in *Arabidopsis* (Huang *et al*., 2026). In line with this observation, we analysed the *AtBRC1* promoter sequence (∼4 kb upstream of the transcription start site) to identify potential TCP4 binding sites. Motif scanning using the JASPAR database identified three putative TCP4 binding motifs. We then reanalysed the ChIP-seq dataset from Dong *et al*. (2019), which revealed a clear binding peak within *AtBRC1* promoter overlapping the first predicted TCP4 binding site (*#1*) (Supplementary Fig. S7).

Based on this, we investigated whether this regulatory mechanism is conserved in rose. We first demonstrated that the expression levels of *RhTCP4a*/*b* and *RhBRC1* in dormant and non-dormant buds from intact or decapitated plants exhibited a clear opposite pattern (Fig. 6A), hinting at a possible inhibition of *RhBRC1* by *RhTCP4a*/*b.* The observation that RhTCP4a and RhTCP4b fusions with GFP are located in the nucleus also supports their roles as regulators of *BRC1* expression (Fig. 6B). Two predicted TCP binding sites were found in the *RhBRC1* promoter (700 and 80 bp upstream of the ATG) (Fig. 6C). Second, to check whether RhTCP4a and RhTCP4b can directly bind to these sites of the *RhBRC1* promoter, we performed an electrophoretic mobility shift assay (EMSA). The results showed that both RhTCP4s proteins bound efficiently to the site 2 of the promoter, with a higher affinity of RhTCP4a compared to RhTCP4b, as confirmed with the addition of mutated competitors (Fig. 6D). In contrast, binding to site 1 appeared weaker and of lower affinity than to site 2 (Supplementary Fig. S8). Finally, to determine whether RhTCP4a and RhTCP4b regulate *RhBRC1* promoter activity, we performed transient co-transfection assays in *Nicotiana benthamiana* leaves. The *RhBRC1* promoter driving the *LUCIFERASE* (*LUC*) reporter was expressed alone or co-expressed with constructs constitutively expressing *RhTCP4a* or *RhTCP4b*. A reduced LUC activity was observed in presence of both *RhTCP4s* constructs, indicating that the activity of the *RhBRC1* promoter is repressed by both TCP4 factors from rose, with repression being more pronounced with RhTCP4a (Fig. 6E). Together, these results provide evidence that RhTCP4a and RhTCP4b directly bind the *RhBRC1* promoter and act as negative regulators of its activity.

**Fig. 6.**
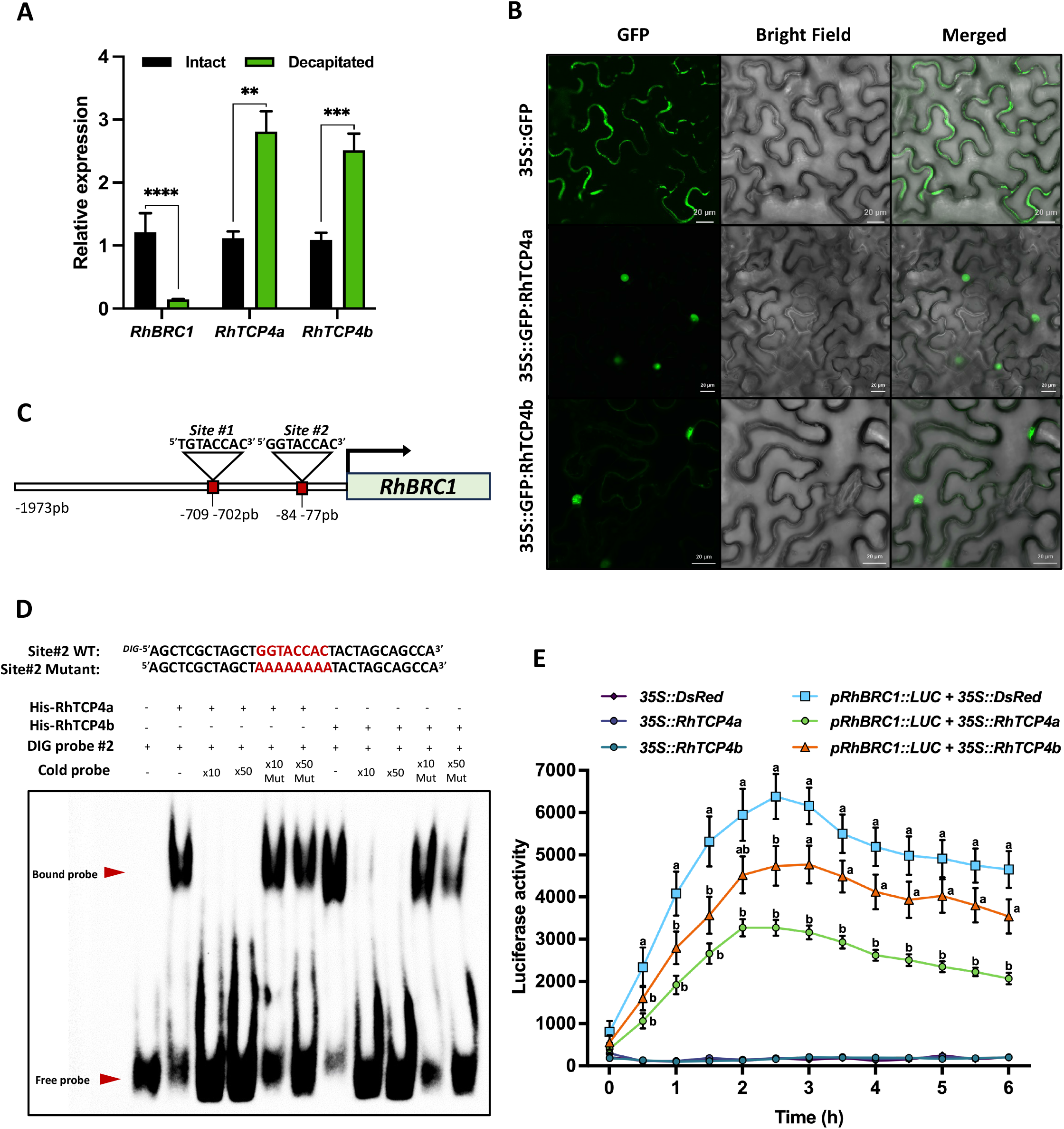
RhTCP4 transcription factor directly represses *RhBRC1* expression. (A) Levels of *RhBRC1* and *RhTCP4a* or *RhTCP4b* in rose axillary buds in intact plants and 24 h after decapitation. (B) Subcellular localization of 35S::GFP:RhTCP4a and 35S::GFP:RhTCP4b in *Nicotiana benthamiana* leaves. (C) Predicted binding sites of RhTCP4s on the *RhBRC1* promoter. (D) EMSA assay using 6xHisTag fused RhTCP4a and RhTCP4b against the site #2 DNA fragments of *RhBRC1* promoter. (E) Luciferase activity levels in *N*. *benthamiana* leaves. The *RhBRC1* promoter fused to the *LUCIFERASE* (*LUC*) reporter gene was co-infiltrated with either RhTCP4a+DsRed, RhTCP4b+DsRed, or DsRed. The effectors 35S::RhTCP4a/b+DsRed were also transfected alone as negative controls. LUC activity was measured every 30 minutes. In (A), data represent the mean ± SD from three biological replicates, and in (E), the mean ± SEM (*n* = 16). Letters indicate significant differences (*P* < 0.05) based on a non-parametric Kruskal–Wallis test followed by Dunn’s post hoc test.

## Discussion

Shoot branching is a fundamental architectural trait governed by endogenous and exogenous cues. Our findings identify the *miR319*–*TCP4* module as a new key transcriptional relay controlling bud activation in response to sugar metabolism via direct repression of *BRC1*. In rose, we propose a model (Fig. 7) in which *RhmiR319* is regulated by carbon metabolism, notably glycolysis/TCA cycle and the OPPP pathway, two metabolic hubs influenced by the antagonistic crosstalk between auxin and sucrose (Wang *et al*., 2021*a*), bud dormancy release (decapitation) and environmental cues such as light intensity. Under high carbon availability conditions, enhanced sugar metabolism is associated with repression of *RhmiR319*, thereby alleviating its inhibition of *RhTCP4a*/*b*. The resulting increase in *RhTCP4a*/*b* levels enables direct binding to the *RhBRC1* promoter, repressing its transcription and ultimately promoting axillary bud outgrowth. In addition, our genetic evidence from *Arabidopsis thaliana* mutants suggests this regulatory module is conserved across dicots. The TCP4/BRC1 regulatory axis acts downstream of SLs (Huang *et al*., 2026), but whether it also integrates sugar metabolic and signalling pathways in *Arabidopsis* remains to be established. Altogether, the *miR319–TCP4* module acts as a conserved molecular integrator, linking local carbon availability to bud growth in rose (Fig. 7).

**Fig. 7.**
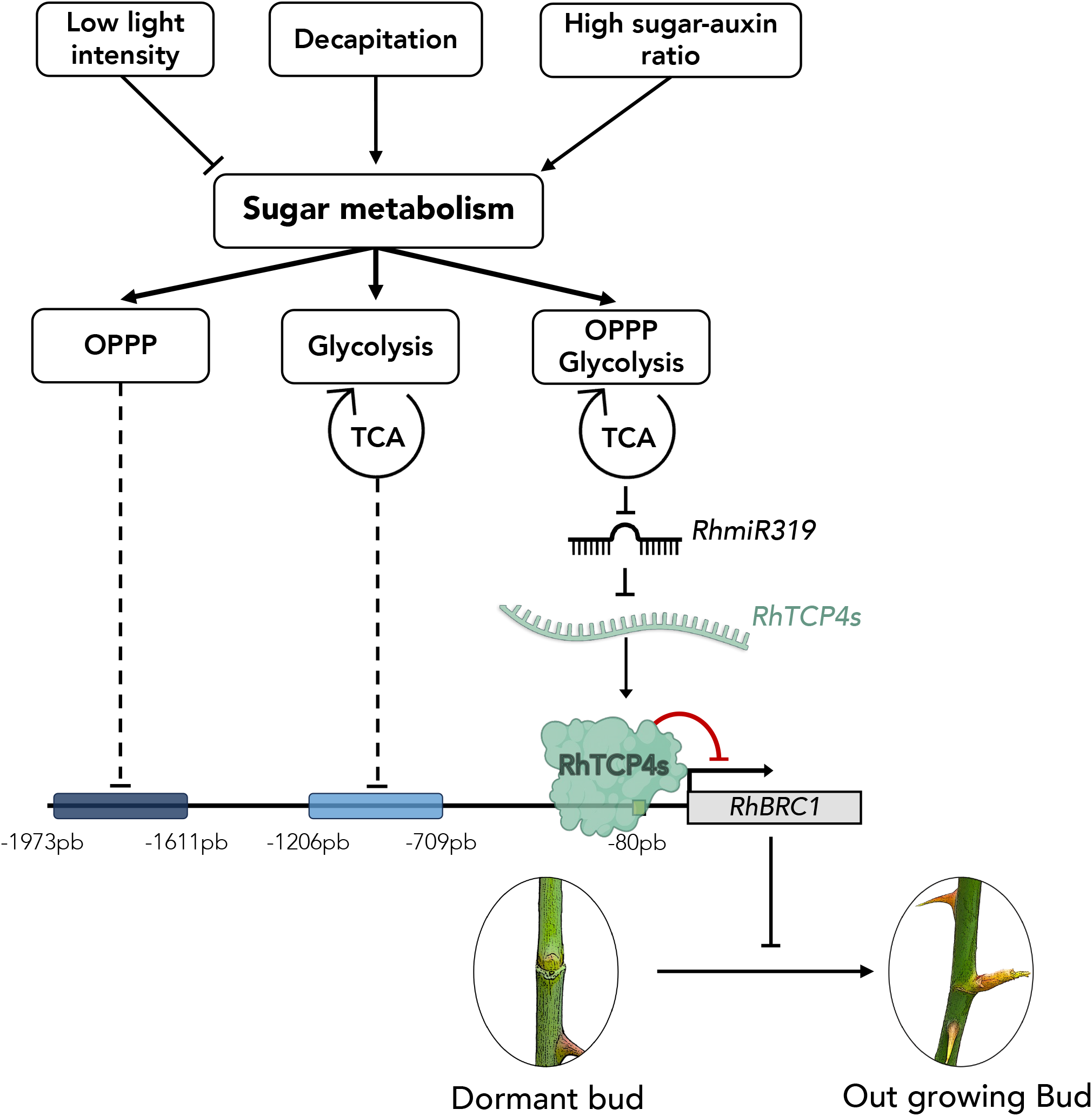
Proposed working model for the role of *RhmiR319*-*RhTCP4s* module in axillary bud outgrowth in *Rosa* in response to sugar metabolic and signalling pathways. Environmental and developmental signals, including light intensity, decapitation and high sugar–auxin ratio, modulate sugar metabolism through the glycolysis/tricarboxylic acid cycle (TCA) and the oxidative pentose phosphate pathways (OPPP). Our previous data showed that these metabolic pathways regulate *RhBRC1* expression *via* two distinct regions of its promoter in rose (Wang *et al*., 2021a). Our results identify an additional pathway in which the combined activity of the glycolysis/TCA cycle and OPPP negatively regulate *RhmiR319*. Reduced *Rhmi319* abundance relieves the repression of *RhTCP4a*/*b,* enabling RhTCP4 proteins to bind a proximal region of the *RhBRC1* promoter, and represses its expression. These findings reveal a new regulatory region of *RhBRC1* linking the *RhmiR319*–*RhTCP4* module to the control of sugar metabolism in shoot branching.

### The *miR319*-*TCP4* module: a new player in the regulation of shoot branching

The *miR319*–*TCP* module has been extensively described for its multiple roles in regulating diverse development processes (Palatnik *et al*., 2003; Schommer *et al*., 2008; Nag *et al*., 2009; Challa *et al*., 2016; Hou *et al*., 2020; Zheng *et al*., 2022; Lan *et al*., 2023; Carvalho *et al*., 2025). In monocots, the *OsmiR319*/*OsTCP21*-*OsGAmyb* or *TamiR319*/*TaGAMYB3* modules regulate tillering, with opposite effects depending on the species (Zhou *et al*., 2013; Wang *et al*., 2021*b*; Jian *et al*., 2022). Here, we extend miR319’s role in branching control to dicot species and link this function to their regulation of evolutionarily conserved targets, the TCP genes. In *Arabidopsis*, elevated levels of *miR319*, whether through endogenous overexpression (*jaw-D*) or heterologous expression of rose-derived *miR319* (*35S::RhmiR319*), consistently reduced lateral branching number (Fig. 4A, B). *miR319* represses five CIN-TCP transcription factors (*TCP2, TCP3, TCP4, TCP10, TCP24*) (Palatnik *et al*., 2003), pointing to a role of one or more of these targets in branching regulation. Because neither the *tcp4* nor the *tcp10* single mutant exhibits any significant branching change, whereas the *tcp4/10* double mutant displays reduced branching, TCP4 and TCP10 appeared as redundant actors promoting *Arabidopsis* shoot branching (Fig. 5A, B and supplementary Fig. S5). In contrast to a recent report (Huang *et al*., 2026), TCP4 is sufficient to promote shoot branching, as expression of *miR319*-insensitive *TCP4* variants (*soj8*, *35S::AtmTCP4*) increases the number of primary rosette branches (Fig. 5D, E). Similarly, overexpression of *miR319*-resistant rose TCP4 orthologs (*RhmTCP4a* and *RhmTCP4b*) in *Arabidopsis* also enhanced branching, indicating that the branch-promoting function of TCP4 is evolutionary conserved (Fig. 5G, H). Strikingly, while most previously described miRNA modules, such as *miR156*–*SPL14* and *miR6288*–*ppTCP4* (peach *BRC2* ortholog), promote branching in dicots, our results indicate that *miR319*–*TCP4* is another inhibitory module as reported previously for *miR171*–*SCL6* (Schwarz *et al*., 2008; Wang *et al*., 2010; Curaba *et al*., 2013; Barrera-Rojas *et al*., 2023; Wang *et al*., 2024).

### Sugar metabolism regulates *RhmiR319*–RhTCP4s during bud outgrowth in roses

Bud outgrowth stimulation is governed by multiple internal and environmental cues, including hormonal, metabolic and light signals (Domagalska and Leyser, 2011; Leduc *et al*., 2014; Mason *et al*., 2014; Barbier *et al*., 2015; Roman *et al*., 2016; Bertheloot *et al*., 2020). Consistent with this, we previously demonstrated that bud outgrowth in roses is regulated by sugar metabolism and related signalling, notably through glycolysis/TCA cycle and the OPPP (Wang *et al*., 2021*a*). However, how sugar metabolism mechanistically acts on bud outgrowth remained largely unresolved. Here, we identified the *RhmiR319–RhTCP4a*/*b* module as a key component linking sugar metabolism and signalling to bud outgrowth. Indeed, combined inhibition of glycolysis/TCA cycle and the OPPP, either directly by inhibitors or indirectly *via* auxin, promotes accumulation of mature *RhmiR319* and downregulation of its targets, *RhTCP4a* and *RhTCP4b* (Fig. 2C, D). This link between sugar metabolism and regulation of the *RhmiR319-TCP4* module is also supported at the plant level. Decapitation, which stimulates sugar metabolism (Wang *et al*., 2021*a*), is accompanied by decreased *RhmiR319* expression and increased *RhTCP4a/b* transcript levels (Fig. 3C). Conversely, under low light conditions limiting carbon availability, the opposite pattern is observed (Fig. 3D).

The positive regulation of *TCP4* by sugar metabolism-related signals (Fig. 2C, D, 3C, D) and by hormones such as gibberellins in tree peony buds (Wang *et al*., 2025*b*), support a role in integrating metabolic and developmental cues to regulate bud outgrowth in perennial species. In addition, *miR319* and its TCP targets respond to diverse environmental signals across species. In particular, *miR319* expression is induced by cold, salinity, drought but repressed by heat (Zhou *et al*., 2013; Joshi *et al*., 2021; Singh *et al*., 2021), and it also contributes to biotic stress responses (Zhao *et al*., 2015; Wu *et al*., 2020; Dong *et al*., 2021). Together with our findings on the sensitivity of rose bud sugar metabolism and *miR319* levels to external signals (such as light intensity, the daily cycle, and decapitation), these results suggest that sugar metabolism acts as a regulatory hub enabling bud to integrate diverse cues and control bud outgrowth through distinct mechanisms. Notably, several studies reported that unfavourable light conditions (darkness), which reduces bud sink strength for sugars (Girault *et al*., 2010; Henry *et al*., 2011), upregulate *RhmiR319* expression (Mallet, 2022). This highlights the importance of future investigations into the role of the sugar metabolism- *miR319* pathway in regulating shoot branching under stress conditions.

### TCP4 represses *BRC1* and reveals a new level of transcriptional regulation of bud outgrowth

Although *BRC1* expression is repressed by sugars (Mason *et al*., 2014; Barbier *et al*., 2015; Wang *et al*., 2021*a*), the precise molecular mechanisms behind this *BRC1* regulation remain to be elucidated. Our results show that overexpression of *miR319* or loss of function of *tcp4*/*10* causes a decrease in the number of lateral branches (Fig. 4D, 5C), concomitant with increased *BRC1* transcription levels. In contrast, TCP4 upregulation repressed *BRC1* expression (Fig. 5F, I). These results suggest that TCP4 promotes branching by repressing *BRC1* expression and are consistent with the ability of AtTCP4 to bind directly to the *AtBRC1* promoter (Supplementary Fig. S7;(Dong *et al*., 2019; Huang *et al*., 2026). Results of co-transfection and EMSA experiments demonstrated that RhTCP4a and RhTCP4b directly bind to the *RhBRC1* promoter and repress its transcriptional activity (Fig. 6D, E), highlighting the functional conservation of the direct TCP4-*BRC1* regulatory interaction between *Arabidopsis* and *Rosa*. More interestingly, our results identify a novel cis-regulatory element at -80bp upstream of the *RhBRC1* ATG that integrates signals from both glycolysis/TCA cycle and OPPP pathways. This site is distinct from previously characterized regions associated with glycolysis/TCA cycle (−1206 to −709 bp) and OPPP-dependent regulation (−1973 to −1611 bp) (Wang *et al*., 2021*a*), revealing a new sugar-responsive mechanism mediated by *RhmiR319*–*RhTCP4a*/*b*, which integrates signals from these two sugar metabolic pathways (Fig. 7). This highlights the concept that the *RhBRC1* promoter functions as a hub that coordinates diverse signals through distinct transcription factor binding sites. Other factors directly regulating *BRC1* promoter activity include *BRI1*-*EMS*-*Suppressor* (BES1), a transcription factor acting downstream of the brassinosteroid (BR) and SL signalling pathways in *Arabidopsis*. BES1 binds directly to the *BRC1* promoter and, by recruiting D53-like SMXL proteins, probably interacting with TCP4, to form a complex that represses *BRC1* expression, thereby promoting axillary bud outgrowth (Hu *et al*., 2020; Huang *et al*., 2026). Conversely, SPL transcription factors such as SPL9–SPL15 in *Arabidopsis* (Xie *et al*., 2020), or SPL16 and SPL23 in poplar (Wei *et al*., 2024), activate *BRC1* transcription by binding to its promoter. Notably, SPL9–SPL15 are targets of *miR156*, revealing that *BRC1* expression can be controlled through another miRNA–target gene regulatory module (Schwarz *et al*., 2008; Wang *et al*., 2008; Wei *et al*., 2012). This regulation of *BRC1* is conserved in monocots, as the transcription factor OsIPA1/OsSPL14 directly binds to the promoter of *TB1* (a *BRC1* ortholog) in rice and positively activates its transcription (Jiao *et al*., 2010; Lu *et al*., 2013). In addition, *CIRCADIAN CLOCK ASSOCIATED1* (*OsCCA1*), which is repressed by sugars, has been shown to directly bind to the promoters of both *OsIPA1* and *OsTB1* in rice, linking circadian regulation to the control of shoot branching (Wang *et al*., 2020*a*). Finally, *NAM3* transcriptionally activates *BRC1*, providing another example of positive transcriptional regulation in axillary bud control in tomato (Wang *et al*., 2025*a*). BRC1 is also post-translationally regulated. For instance, CUC2, targeted by *miR164*, directly activates BRC1 through protein interaction in cotton (Zhan *et al*., 2021), while TCP14/15 (Gastaldi *et al*., 2024) and TCP Interacting containing EAR motif protein 1 (TIE1) (Yang *et al*., 2018) repress it in *Arabidopsis*. While the *miR156*–*SPL*–*BRC1* cascade directly activates *BRC1* transcription, our results reveal that the *miR319*–*TCP4* module acts as a novel direct repressor of *BRC1*, establishing a complementary mechanism for adjusting expression of this key regulator of shoot branching.

Our study unveils the *miR319*-*TCP4-BRC1* module as a pivotal new link between sugar metabolism and signalling and the control of bud outgrowth (Fig. 7). Future work should define the molecular bridge connecting sugar availability to *miR319*/*TCP4* regulation. Prime candidates are the energy-sensing *SnRK1* kinase (activated under low energy/sugar conditions to reprogram metabolism and promote stress tolerance) and *TOR* kinase (sensing nutrients including sugar availability to regulate growth). It will also be important to identify the transcription factors that tie *BRC1* expression to the glycolysis/TCA cycle or the OPPP (Fig. 7). Investigating how *miR319* and TCP4 responses to sugars-hormones crosstalk, particularly to CKs and SLs, and performing DAP-seq to map RhTCP4 targets will illuminate its broad role in plant architecture. Ultimately, dissecting the *miRNA*-*TCP4* dynamics under environmental stresses, such as drought and heat stress will reveal the contribution of this module to developmental plasticity.

## Supplementary data

**Fig. S1. Conservation of TCP domain truncation of rose TCP proteins. Supports Fig. 2.**

**Fig. S2. *RhmiR319* targets *RhTCP4* genes. Supports Fig. 2.**

**Fig. S3. *RhmiR319* negatively regulates *RhTCP4* genes. Supports Fig. 3.**

**Fig. S4. Leaf morphology of WT*, jaw-D and 35S::RhmiR319* in *Arabidopsis.* Supports Fig. 4.**

**Fig. S5. Shoot branching in *tcp10* line in *Arabidopsis*. Supports Fig. 5.**

**Fig. S6. Branching phenotype of independent *35S::RhmTCP4a/b* lines in *Arabidopsis*. Supports Fig. 5.**

**Fig. S7. TCP4 binds *BRC1* promoter in *Arabidopsis*. Supports Fig. 6.**

**Fig. S8. EMSA assay using 6xHis-tagged RhTCP4a and RhTCP4b proteins with the site #1 DNA fragment of *RhBRC1* promoter. Supports Fig. 6.**

**Methods S1. ChIP-seq data analysis**

**Table S1. Primers list used in this study.**

**Table S2. Primers list for qRT-PCR.**

## Acknowledgements (102 words)

The authors would like to thank Carla Schommer, Tomotsugu Koyama and Javier Palatnik for providing the *jaw-D*, *soj8*, *mTCP4*, *tcp4* and *tcp4*/*10* lines. The authors also thank Maël Baudin for kindly providing the plasmids used to produce the recombinant proteins. We also thank the SFR Quasav for access to the IMAC technical facility for confocal microscopy, the ANAN platform for RNA and DNA quantification and plate reader facilities and PHENOTIC for plant growth facilities used for *Arabidopsis thaliana* and rose experiments. We also thank Kaat Hellyn, Bénédicte Dubuc and Denis Cesbron for their assistance with rose plant multiplication and greenhouse chamber management.

## Author contributions

LG, JLG, SS and PL conceived and designed the research. LG performed all the experiments with help from JL, AP, TP, LO and MDPG. LG and JM designed and performed genetic transformation of the *35S::RhmiR319a* line (JM) and the *35S::RhmTCP4a* and *35S::RhmTCPb* lines (LG). JK performed de novo analysis of the ChIP-seq data. LG wrote the original draft of the paper. LG, JM, JLG, SS, PL, AP, FB revised and edited this manuscript. All authors have read and agreed on the final version.

## Data availability

Raw small RNA sequencing data generated in this study have been deposited in the NCBI Sequence Read Archive (SRA) under BioProject accession PRJNA1470483 (SRA study accession SRP703594).

## References

Aguilar-Martínez José Antonio, Poza-Carrión César, Cubas Pilar. 2007. Arabidopsis BRANCHED1 Acts as an Integrator of Branching Signals within Axillary Buds. The Plant Cell 19, 458–472.

Barbier Francois F, Cao Da, Fichtner Franziska, Weiste Christoph, Perez-Garcia Maria-Dolores, Caradeuc Mathieu, Le Gourrierec José, Sakr Soulaiman, Beveridge CA. 2021. HEXOKINASE1 signalling promotes shoot branching and interacts with cytokinin and strigolactone pathways. New Phytologist 231, 1088–1104.

Barbier FF, Chabikwa TG, Ahsan MU, Cook SE, Powell R, Tanurdzic M, Beveridge CA. 2019. A phenol/chloroform-free method to extract nucleic acids from recalcitrant, woody tropical species for gene expression and sequencing. Plant Methods 15, 62.

Barbier F, Péron T, Lecerf M, et al. 2015. Sucrose is an early modulator of the key hormonal mechanisms controlling bud outgrowth in Rosa hybrida. Journal of Experimental Botany 66, 2569–2582.

Barrera-Rojas CH, Vicente MH, Pinheiro Brito DA, et al. 2023. Tomato miR156-targeted SlSBP15 represses shoot branching by modulating hormone dynamics and interacting with GOBLET and BRANCHED1b. (U Vijayraghavan, Ed.). Journal of Experimental Botany 74, 5124–5139.

Bertheloot J, Barbier F, Boudon F, Perez-Garcia MD, Péron T, Citerne S, Dun E, Beveridge C, Godin C, Sakr S. 2020. Sugar availability suppresses the auxin-induced strigolactone pathway to promote bud outgrowth. New Phytologist 225, 866–879.

Beveridge CA, Rameau C, Wijerathna-Yapa A. 2023. Lessons from a century of apical dominance research. (J Lunn, Ed.). Journal of Experimental Botany 74, 3903–3922.

Boumaza R, Huché-Thélier L, Demotes-Mainard S, et al. 2010. Sensory profiles and preference analysis in ornamental horticulture: The case of the rosebush. Food Quality and Preference 21, 987–997.

Cao D, Chabikwa T, Barbier F, Dun EA, Fichtner F, Dong L, Kerr SC, Beveridge CA. 2023. Auxin-independent effects of apical dominance induce changes in phytohormones correlated with bud outgrowth. Plant Physiology 192, 1420–1434.

Cao J-F, Zhao B, Huang C-C, et al. 2020. The miR319-Targeted GhTCP4 Promotes the Transition from Cell Elongation to Wall Thickening in Cotton Fiber. Molecular Plant 13, 1063–1077.

Carvalho A, Vicente MH, Ferigolo LF, et al. 2025. The MIR319 -based repression of SLTCP2 / LANCEOLATE activity is required for regulating tomato fruit shape. The Plant Journal 121, e17174.

Challa KR, Aggarwal P, Nath U. 2016. Activation of YUCCA5 by the Transcription Factor TCP4 Integrates Developmental and Environmental Signals to Promote Hypocotyl Elongation in Arabidopsis. The Plant Cell 28, 2117–2130.

Challa KR, Rath M, Nath U. 2019. The CIN-TCP transcription factors promote commitment to differentiation in Arabidopsis leaf pavement cells via both auxin-dependent and independent pathways. (L-J Qu, Ed.). PLOS Genetics 15, e1007988.

Clough SJ, Bent AF. 1998. Floral dip: a simplified method for Agrobacterium-mediated transformation of Arabidopsis thaliana. The Plant Journal 16, 735–743.

Curaba J, Talbot M, Li Z, Helliwell C. 2013. Over-expression of microRNA171 affects phase transitions and floral meristem determinancy in barley. BMC Plant Biology 13, 6.

Dai X, Zhuang Z, Zhao PX. 2018. psRNATarget: a plant small RNA target analysis server (2017 release). Nucleic Acids Research 46, W49–W54.

Doidy J, Wang Y, Gouaille L, Goma-Louamba I, Jiang Z, Pourtau N, Le Gourrierec J, Sakr S. 2024. Sugar Transport and Signaling in Shoot Branching. International Journal of Molecular Sciences 25, 13214.

Domagalska MA, Leyser O. 2011. Signal integration in the control of shoot branching. Nature Reviews Molecular Cell Biology 12, 211–221.

Dong W, Ren W, Wang X, Mao Y, He Y. 2021. MicroRNA319a regulates plant resistance to Sclerotinia stem rot. (S Spoel, Ed.). Journal of Experimental Botany 72, 3540–3553.

Dong J, Sun N, Yang J, et al. 2019. The Transcription Factors TCP4 and PIF3 Antagonistically Regulate Organ-Specific Light Induction of SAUR Genes to Modulate Cotyledon Opening during De-Etiolation in Arabidopsis. The Plant Cell 31, 1155–1170.

Dong H, Wang J, Song X, et al. 2023. HY5 functions as a systemic signal by integrating BRC1-dependent hormone signaling in tomato bud outgrowth. Proceedings of the National Academy of Sciences 120, e2301879120.

Duarte GT, Matiolli CC, Pant BD, Schlereth A, Scheible W-R, Stitt M, Vicentini R, Vincentz M. 2013. Involvement of microRNA-related regulatory pathways in the glucose-mediated control of Arabidopsis early seedling development. Journal of Experimental Botany 64, 4301–4312.

Dun EA, De Saint Germain A, Rameau C, Beveridge CA. 2012. Antagonistic Action of Strigolactone and Cytokinin in Bud Outgrowth Control. Plant Physiology 158, 487–498.

Efroni I, Blum E, Goldshmidt A, Eshed Y. 2008. A Protracted and Dynamic Maturation Schedule Underlies Arabidopsis Leaf Development. The Plant Cell 20, 2293–2306.

Fan D, Ran L, Hu J, Ye X, Xu D, Li J, Su H, Wang X, Ren S, Luo K. 2020. miR319a/TCP module and DELLA protein regulate trichome initiation synergistically and improve insect defenses in Populus tomentosa. New Phytologist 227, 867–883.

Fang Y, Zheng Y, Lu W, Li J, Duan Y, Zhang S, Wang Y. 2021. Roles of miR319-regulated TCPs in plant development and response to abiotic stress. The Crop Journal 9, 17–28.

Fichtner F, Barbier FF, Annunziata MG, Feil R, Olas JJ, Mueller-Roeber B, Stitt M, Beveridge CA, Lunn JE. 2021a. Regulation of shoot branching in arabidopsis by trehalose 6-phosphate. New Phytologist 229, 2135–2151.

Fichtner F, Barbier FF, Annunziata MG, Feil R, Olas JJ, Mueller-Roeber B, Stitt M, Beveridge CA, Lunn JE. 2021b. Regulation of shoot branching in arabidopsis by trehalose 6-phosphate. New Phytologist 229, 2135–2151.

Fichtner F, Barbier FF, Feil R, Watanabe M, Annunziata MG, Chabikwa TG, Höfgen R, Stitt M, Beveridge CA, Lunn JE. 2017. Trehalose 6-phosphate is involved in triggering axillary bud outgrowth in garden pea (Pisum sativum L.). The Plant Journal 92, 611–623.

Fichtner F, Humphreys JL, Barbier FF, Feil R, Westhoff P, Moseler A, Lunn JE, Smith SM, Beveridge CA. 2024. Strigolactone signalling inhibits trehalose 6-phosphate signalling independently of BRC1 to suppress shoot branching. New Phytologist 244, 900–913.

Fu C, Sunkar R, Zhou C, et al. 2012. Overexpression of miR156 in switchgrass (Panicum virgatum L.) results in various morphological alterations and leads to improved biomass production. Plant Biotechnology Journal 10, 443–452.

Gälweiler L, Guan C, Müller A, Wisman E, Mendgen K, Yephremov A, Palme K. 1998. Regulation of Polar Auxin Transport by AtPIN1 in Arabidopsis Vascular Tissue. Science 282, 2226–2230.

Gastaldi V, Nicolas M, Muñoz-Gasca A, Cubas P, Gonzalez DH, Lucero L. 2024. Class I TCP transcription factors TCP14 and TCP15 promote axillary branching in Arabidopsis by counteracting the action of Class II TCP BRANCHED1. New Phytologist 243, 1810–1822.

Girault T, Abidi F, Sigogne M, Pelleschi-Travier S, Boumaza R, Sakr S, Leduc N. 2010. Sugars are under light control during bud burst in Rosa sp.: Photocontrol of sugars during bud burst. Plant, Cell & Environment, no-no.

Göbel M, Fichtner F. 2023. Functions of sucrose and trehalose 6-phosphate in controlling plant development. Journal of Plant Physiology 291, 154140.

González-Grandío E, Pajoro A, Franco-Zorrilla JM, Tarancón C, Immink RGH, Cubas P. 2017. Abscisic acid signaling is controlled by a BRANCHED1/HD-ZIP I cascade in Arabidopsis axillary buds. Proceedings of the National Academy of Sciences 114.

Guo S, Xu Y, Liu H, Mao Z, Zhang C, Ma Y, Zhang Q, Meng Z, Chong K. 2013. The interaction between OsMADS57 and OsTB1 modulates rice tillering via DWARF14. Nature Communications 4, 1566.

Hallé F, Oldeman RA. 1970. Essai sur l’architecture et la dynamique de croissance des arbres tropicaux.

Henry C, Rabot A, Laloi M, Mortreau E, Sigogne M, Leduc N, Lemoine R, Sakr S, Vian A, Pelleschi-Travier S. 2011. Regulation of RhSUC2, a sucrose transporter, is correlated with the light control of bud burst in Rosa sp.: Sucrose transporter role in bud burst. Plant, Cell & Environment 34, 1776–1789.

Hibrand Saint-Oyant L, Ruttink T, Hamama L, et al. 2018. A high-quality genome sequence of Rosa chinensis to elucidate ornamental traits. Nature Plants 4, 473–484.

Hou J, Xu H, Fan D, Ran L, Li J, Wu S, Luo K, He X. 2020. MiR319a-targeted PtoTCP20 regulates secondary growth via interactions with PtoWOX4 and PtoWND6 in Populus tomentosa. New Phytologist 228, 1354–1368.

Hu J, Ji Y, Hu X, Sun S, Wang X. 2020. BES1 Functions as the Co-regulator of D53-like SMXLs to Inhibit BRC1 Expression in Strigolactone-Regulated Shoot Branching in Arabidopsis. Plant Communications 1, 100014.

Huang Y, Bi L, Huang Y, Liu J, Wang L, Qiu F, Wang Y, Chen L, Zhang M, Yao R. 2026. A group of TCP transcription factors is a missing link in strigolactone signaling. Journal of Integrative Plant Biology, jipb.70281.

Huang J, Li Z, Zhao D. 2016. Deregulation of the OsmiR160 Target Gene OsARF18 Causes Growth and Developmental Defects with an Alteration of Auxin Signaling in Rice. Scientific Reports 6, 29938.

Jang JC, Sheen J. 1994. Sugar sensing in higher plants. The Plant Cell 6, 1665–1679.

Jian C, Hao P, Hao C, et al. 2022. The miR319/TaGAMYB3 module regulates plant architecture and improves grain yield in common wheat (Triticum aestivum). New Phytologist 235, 1515–1530.

Jiang Z, Wang M, Gouaille L, et al. 2025. Cytokinin-induced bud outgrowth depends on sugar metabolism and signalling. (R Napier, Ed.). Journal of Experimental Botany 76, 5351–5366.

Jiao Y, Wang Y, Xue D, et al. 2010. Regulation of OsSPL14 by OsmiR156 defines ideal plant architecture in rice. Nature Genetics 42, 541–544.

Johnson NR, Yeoh JM, Coruh C, Axtell MJ. 2016. Improved Placement of Multi-mapping Small RNAs. G3 Genes|Genomes|Genetics 6, 2103–2111.

Jones DT, Taylor WR, Thornton JM. 1992. The rapid generation of mutation data matrices from protein sequences. Bioinformatics 8, 275–282.

Joshi GAN, Chauhan C, Das S. 2021. Sequence and functional analysis of MIR319 promoter homologs from Brassica juncea reveals regulatory diversification and altered expression under stress. Molecular Genetics and Genomics 296, 731–749.

Kebrom TH, Chandler PM, Swain SM, King RW, Richards RA, Spielmeyer W. 2012. Inhibition of Tiller Bud Outgrowth in the tin Mutant of Wheat Is Associated with Precocious Internode Development. Plant Physiology 160, 308–318.

Kebrom TH, Mullet JE. 2015. Photosynthetic leaf area modulates tiller bud outgrowth in sorghum. Plant, Cell & Environment 38, 1471–1478.

Köhler E, Barrach H-J, Neubert D. 1970. Inhibition of NADP dependent oxidoreductases by the 6-aminonicotinamide analogue of NADP. FEBS Letters 6, 225–228.

Koyama T, Furutani M, Tasaka M, Ohme-Takagi M. 2007. TCP Transcription Factors Control the Morphology of Shoot Lateral Organs via Negative Regulation of the Expression of Boundary-Specific Genes in Arabidopsis. The Plant Cell 19, 473–484.

Koyama T, Kunieda T, Toyonaga H, et al. 2025. TCP3 -mediated regulation of cell expansion in Arabidopsis thaliana. New Phytologist 248, 2981–2995.

Koyama T, Sato F, Ohme-Takagi M. 2017. Roles of miR319 and TCP Transcription Factors in Leaf Development. Plant Physiology 175, 874–885.

Kozomara A, Birgaoanu M, Griffiths-Jones S. 2019. miRBase: from microRNA sequences to function. Nucleic Acids Research 47, D155–D162.

Kravchik M, Stav R, Belausov E, Arazi T. 2019. Functional Characterization of microRNA171 Family in Tomato. Plants 8, 10.

Kumar S, Stecher G, Suleski M, Sanderford M, Sharma S, Tamura K. 2024. MEGA12: Molecular Evolutionary Genetic Analysis Version 12 for Adaptive and Green Computing. (FU Battistuzzi, Ed.). Molecular Biology and Evolution 41, msae263.

Lam HM, Peng Ssy, Coruzzi GM. 1994. Metabolic Regulation of the Gene Encoding Glutamine-Dependent Asparagine Synthetase in Arabidopsis thaliana. Plant Physiology 106, 1347–1357.

Lan J, Wang N, Wang Y, Jiang Y, Yu H, Cao X, Qin G. 2023. Arabidopsis TCP4 transcription factor inhibits high temperature-induced homeotic conversion of ovules. Nature Communications 14, 5673.

Langmead B, Trapnell C, Pop M, Salzberg SL. 2009. Ultrafast and memory-efficient alignment of short DNA sequences to the human genome. Genome Biology 10, R25.

Leduc N, Roman H, Barbier F, Péron T, Huché-Thélier L, Lothier J, Demotes-Mainard S, Sakr S. 2014. Light Signaling in Bud Outgrowth and Branching in Plants. Plants 3, 223–250.

Lejay L, Gansel X, Cerezo M, Tillard P, Müller C, Krapp A, von Wirén N, Daniel-Vedele F, Gojon A. 2003. Regulation of Root Ion Transporters by Photosynthesis: Functional Importance and Relation with Hexokinase. The Plant Cell 15, 2218–2232.

Lejay L, Wirth J, Pervent M, Cross JM-F, Tillard P, Gojon A. 2008. Oxidative Pentose Phosphate Pathway-Dependent Sugar Sensing as a Mechanism for Regulation of Root Ion Transporters by Photosynthesis. Plant Physiology 146, 2036–2053.

Li H, Handsaker B, Wysoker A, Fennell T, Ruan J, Homer N, Marth G, Abecasis G, Durbin R, 1000 Genome Project Data Processing Subgroup. 2009. The Sequence Alignment/Map format and SAMtools. Bioinformatics 25, 2078–2079.

Li Y, He Y, Qin T, Guo X, Xu K, Xu C, Yuan W. 2023. Functional conservation and divergence of miR156 and miR529 during rice development. The Crop Journal 11, 692–703.

Li M, Li H, Zhu Q, Liu D, Li Z, Chen H, Luo J, Gong P, Ismail AM, Zhang Z. 2024. Knockout of the sugar transporter OSSTP15 enhances grain yield by improving tiller number due to increased sugar content in the shoot base of rice (Oryza sativa L.). New Phytologist 241, 1250–1265.

Li X, Xia K, Liang Z, Chen K, Gao C, Zhang M. 2016. MicroRNA393 is involved in nitrogen-promoted rice tillering through regulation of auxin signal transduction in axillary buds. Scientific Reports 6, 32158.

Lian H, Wang L, Ma N, Zhou C-M, Han L, Zhang T-Q, Wang J-W. 2021. Redundant and specific roles of individual MIR172 genes in plant development. (X Chen, Ed.). PLOS Biology 19, e3001044.

Liu Y, Chen S, Pal S, Yu J, Zhou Y, Tran L-SP, Xia X. 2024. The hormonal, metabolic, and environmental regulation of plant shoot branching. New Crops 1, 100028.

Liu J, Cheng X, Liu P, Sun J. 2017. miR156-Targeted SBP-Box Transcription Factors Interact with DWARF53 to Regulate TEOSINTE BRANCHED1 and BARREN STALK1 Expression in Bread Wheat. Plant Physiology 174, 1931–1948.

Liu S, Mi X, Zhang R, An Y, Zhou Q, Yang T, Xia X, Guo R, Wang X, Wei C. 2019a. Integrated analysis of miRNAs and their targets reveals that miR319c/TCP2 regulates apical bud burst in tea plant (Camellia sinensis). Planta 250, 1111–1129.

Liu W, Peng B, Song A, Jiang J, Chen F. 2019b. Sugar Transporter, CmSWEET17, Promotes Bud Outgrowth in Chrysanthemum Morifolium. Genes 11, 26.

Ljung K, Bhalerao RP, Sandberg G. 2002. Sites and homeostatic control of auxin biosynthesis in Arabidopsis during vegetative growth: Auxin biosynthesis and distribution in Arabidopsis. The Plant Journal 28, 465–474.

Lopez-Obando M, Ligerot Y, Bonhomme S, Boyer F-D, Rameau C. 2015. Strigolactone biosynthesis and signaling in plant development. Development 142, 3615–3619.

Lu Z, Yu H, Xiong G, et al. 2013. Genome-Wide Binding Analysis of the Transcription Activator IDEAL PLANT ARCHITECTURE1 Reveals a Complex Network Regulating Rice Plant Architecture. The Plant Cell 25, 3743–3759.

Mallet J. 2022. Régulations post-transcriptionnelles par les miARNs du débourrement des bourgeons et de son photo-contrôle chez le rosier buisson Rosa ‘Radrazz’. Doctorat ès Sciences agronomiques, PhD, Angers.

Mallet J, Laufs P, Leduc N, Le Gourrierec J. 2022. Photocontrol of Axillary Bud Outgrowth by MicroRNAs: Current State-of-the-Art and Novel Perspectives Gained From the Rosebush Model. Frontiers in Plant Science 12, 770363.

Mammarella MF, Lucero L, Hussain N, et al. 2023. Long noncoding RNA-mediated epigenetic regulation of auxin-related genes controls shade avoidance syndrome in Arabidopsis. The EMBO Journal 42, EMBJ2023113941.

Martín-Fontecha ES, Tarancón C, Cubas P. 2018. To grow or not to grow, a power-saving program induced in dormant buds. Current Opinion in Plant Biology 41, 102–109.

Mason MG, Ross JJ, Babst BA, Wienclaw BN, Beveridge CA. 2014. Sugar demand, not auxin, is the initial regulator of apical dominance. Proceedings of the National Academy of Sciences 111, 6092–6097.

Mathan J, Bhattacharya J, Ranjan A. 2016. Enhancing crop yield by optimizing plant developmental features. Development 143, 3283–3294.

May P, Liao W, Wu Y, Shuai B, Richard McCombie W, Zhang MQ, Liu QA. 2013. The effects of carbon dioxide and temperature on microRNA expression in Arabidopsis development. Nature Communications 4, 2145.

Morea EGO, Da Silva EM, E Silva GFF, Valente GT, Barrera Rojas CH, Vincentz M, Nogueira FTS. 2016. Functional and evolutionary analyses of the miR156 and miR529 families in land plants. BMC Plant Biology 16, 40.

Nag A, King S, Jack T. 2009. miR319a targeting of TCP4 is critical for petal growth and development in Arabidopsis. Proceedings of the National Academy of Sciences 106, 22534–22539.

Ori N, Cohen AR, Etzioni A, et al. 2007. Regulation of LANCEOLATE by miR319 is required for compound-leaf development in tomato. Nature Genetics 39, 787–791.

Otori K, Tamoi M, Tanabe N, Shigeoka S. 2017. Enhancements in sucrose biosynthesis capacity affect shoot branching in Arabidopsis. Bioscience, Biotechnology, and Biochemistry 81, 1470–1477.

Palatnik JF, Allen E, Wu X, Schommer C, Schwab R, Carrington JC, Weigel D. 2003. Control of leaf morphogenesis by microRNAs. Nature 425, 257–263.

Palatnik JF, Wollmann H, Schommer C, et al. 2007. Sequence and Expression Differences Underlie Functional Specialization of Arabidopsis MicroRNAs miR159 and miR319. Developmental Cell 13, 115– 125.

Porcher A, Guérin V, Leduc N, Lebrec A, Lothier J, Vian A. 2021. Ascorbate–glutathione pathways mediated by cytokinin regulate H2O2 levels in light-controlled rose bud burst. Plant Physiology 186, 910– 928.

Porcher A, Guérin V, Montrichard F, Lebrec A, Lothier J, Vian A. 2020. Ascorbate glutathione-dependent H2O2 scavenging is an important process in axillary bud outgrowth in rosebush. Annals of Botany 126, 1049–1062.

Rabot A, Henry C, Ben Baaziz K, et al. 2012. Insight into the Role of Sugars in Bud Burst Under Light in the Rose. Plant and Cell Physiology 53, 1068–1082.

Rabot A, Portemer V, Péron T, Mortreau E, Leduc N, Hamama L, Coutos-Thévenot P, Atanassova R, Sakr S, Le Gourrierec J. 2014. Interplay of Sugar, Light and Gibberellins in Expression of Rosa hybrida Vacuolar Invertase 1 Regulation. Plant and Cell Physiology 55, 1734–1748.

Rameau C, Bertheloot J, Leduc N, Andrieu B, Foucher F, Sakr S. 2015. Multiple pathways regulate shoot branching. Frontiers in Plant Science 5.

Rodriguez RE, Mecchia MA, Debernardi JM, Schommer C, Weigel D, Palatnik JF. 2010. Control of cell proliferation in Arabidopsis thaliana by microRNA miR396. Development 137, 103–112.

Roman H, Girault T, Barbier F, et al. 2016. Cytokinins Are Initial Targets of Light in the Control of Bud Outgrowth. Plant Physiology 172, 489–509.

Sakr S, Wang M, Dédaldéchamp F, Perez-Garcia M-D, Ogé L, Hamama L, Atanassova R. 2018. The Sugar-Signaling Hub: Overview of Regulators and Interaction with the Hormonal and Metabolic Network. International Journal of Molecular Sciences 19, 2506.

Sarrion-Perdigones A, Falconi EE, Zandalinas SI, Juárez P, Fernández-del-Carmen A, Granell A, Orzaez D. 2011. GoldenBraid: An Iterative Cloning System for Standardized Assembly of Reusable Genetic Modules. (J Peccoud, Ed.). PLoS ONE 6, e21622.

Sarrion-Perdigones A, Vazquez-Vilar M, Palaci J, Castelijns B, Forment J, Ziarsolo P, Blanca J, Granell A, Orzaez D. 2013. GoldenBraid 2.0: A Comprehensive DNA Assembly Framework for Plant Synthetic Biology. PLANT PHYSIOLOGY 162, 1618–1631.

Sarvepalli K, Nath U. 2011. Hyper-activation of the TCP4 transcription factor in Arabidopsis thaliana accelerates multiple aspects of plant maturation. The Plant Journal 67, 595–607.

Schommer C, Debernardi JM, Bresso EG, Rodriguez RE, Palatnik JF. 2014. Repression of Cell Proliferation by miR319-Regulated TCP4. Molecular Plant 7, 1533–1544.

Schommer C, Palatnik JF, Aggarwal P, Chételat A, Cubas P, Farmer EE, Nath U, Weigel D. 2008. Control of Jasmonate Biosynthesis and Senescence by miR319 Targets. (JC Carrington, Ed.). PLoS Biology 6, e230.

Schwarz S, Grande AV, Bujdoso N, Saedler H, Huijser P. 2008. The microRNA regulated SBP-box genes SPL9 and SPL15 control shoot maturation in Arabidopsis. Plant Molecular Biology 67, 183–195.

Shankar N, Sunkara P, Nath U. 2023. A double-negative feedback loop between miR319c and JAW-TCPs establishes growth pattern in incipient leaf primordia in Arabidopsis thaliana. (S Hake, Ed.). PLOS Genetics 19, e1010978.

Singh P, Dutta P, Chakrabarty D. 2021. miRNAs play critical roles in response to abiotic stress by modulating cross-talk of phytohormone signaling. Plant Cell Reports 40, 1617–1630.

Snow R. 1929. The Transmission of Inhibition through Dead Stretches of Stem. Annals of Botany os-43, 261–267.

Stitz M, Kuster D, Reinert M, et al. 2023. TOR acts as a metabolic gatekeeper for auxin-dependent lateral root initiation in Arabidopsis thaliana. The EMBO Journal 42, EMBJ2022111273.

Sun Z, Su C, Yun J, et al. 2019. Genetic improvement of the shoot architecture and yield in soya bean plants via the manipulation of GmmiR156b. Plant Biotechnology Journal 17, 50–62.

Sun X, Wang C, Xiang N, Li X, Yang S, Du J, Yang Y, Yang Y. 2017. Activation of secondary cell wall biosynthesis by miR319-targeted TCP4 transcription factor. Plant Biotechnology Journal 15, 1284–1294.

Thimann KV, Skoog F. 1933. Studies on the Growth Hormone of Plants: III. The Inhibiting Action of the Growth Substance on Bud Development. Proceedings of the National Academy of Sciences 19, 714–716.

Vadde BVL, Challa KR, Nath U. 2018. The TCP 4 transcription factor regulates trichome cell differentiation by directly activating GLABROUS INFLORESCENCE STEMS in Arabidopsis thaliana. The Plant Journal 93, 259–269.

Van Es SW, Muñoz-Gasca A, Romero-Campero FJ, et al. 2024. A gene regulatory network critical for axillary bud dormancy directly controlled by Arabidopsis BRANCHED1. New Phytologist 241, 1193–1209.

Varet H, Brillet-Guéguen L, Coppée J-Y, Dillies M-A. 2016. SARTools: A DESeq2- and EdgeR-Based R Pipeline for Comprehensive Differential Analysis of RNA-Seq Data. (K Mills, Ed.). PLOS ONE 11, e0157022.

Varkonyi-Gasic E, Wu R, Wood M, Walton EF, Hellens RP. 2007. Protocol: a highly sensitive RT-PCR method for detection and quantification of microRNAs. Plant Methods 3, 12.

Wahl V, Ponnu J, Schlereth A, Arrivault S, Langenecker T, Franke A, Feil R, Lunn JE, Stitt M, Schmid M. 2013. Regulation of Flowering by Trehalose-6-Phosphate Signaling in Arabidopsis thaliana. Science 339, 704–707.

Wang F, Han T, Song Q, Ye W, Song X, Chu J, Li J, Chen ZJ. 2020a. The Rice Circadian Clock Regulates Tiller Growth and Panicle Development Through Strigolactone Signaling and Sugar Sensing. The Plant Cell 32, 3124–3138.

Wang M, Le Moigne M-A, Bertheloot J, Crespel L, Perez-Garcia M-D, Ogé L, Demotes-Mainard S, Hamama L, Davière J-M, Sakr S. 2019a. BRANCHED1: A Key Hub of Shoot Branching. Frontiers in Plant Science 10, 76.

Wang J, Li J, Dong Y, Dong H, Zhou J, Shi K, Zhou Y, Yu J, Xia X. 2025a. CIPK1 -regulated transcription factor NAM3 coordinates shoot branching and nitrate accumulation in tomato. New Phytologist 248, 1940–1958.

Wang Q, Li B, Qiu Z, Ying J, Jin X, Lu Z, Zhang J, Chen X, Zhu X. 2025b. The involvement of PsTCP genes in hormone-mediated process of bud dormancy release in tree peony (Paeonia suffruticosa). BMC Genomics 26, 266.

Wang L, Mai Y-X, Zhang Y-C, Luo Q, Yang H-Q. 2010. MicroRNA171c-Targeted SCL6-II, SCL6-III, and SCL6-IV Genes Regulate Shoot Branching in Arabidopsis. Molecular Plant 3, 794–806.

Wang T, Miao M, Zhao J, Kumar A, Li X. 2025c. Sugars Integrate External and Internal Signals in Regulating Shoot Branching. Plant, Cell & Environment 48, 8688–8701.

Wang M, Ogé L, Voisine L, Perez-Garcia M-D, Jeauffre J, Hibrand Saint-Oyant L, Grappin P, Hamama L, Sakr S. 2019b. Posttranscriptional Regulation of RhBRC1 (Rosa hybrida BRANCHED1) in Response to Sugars is Mediated via its Own 3′ Untranslated Region, with a Potential Role of RhPUF4 (Pumilio RNA-Binding Protein Family). International Journal of Molecular Sciences 20, 3808.

Wang M, Pérez-Garcia M-D, Davière J-M, et al. 2021a. Outgrowth of the axillary bud in rose is controlled by sugar metabolism and signalling. (P Cubas, Ed.). Journal of Experimental Botany 72, 3044–3060.

Wang J-W, Schwab R, Czech B, Mica E, Weigel D. 2008. Dual Effects of miR156-Targeted SPL Genes and CYP78A5/KLUH on Plastochron Length and Organ Size in Arabidopsis thaliana. The Plant Cell 20, 1231–1243.

Wang H, Wang H. 2015. The miR156/SPL Module, a Regulatory Hub and Versatile Toolbox, Gears up Crops for Enhanced Agronomic Traits. Molecular Plant 8, 677–688.

Wang Y, Wang Z, Amyot L, Tian L, Xu Z, Gruber MY, Hannoufa A. 2015. Ectopic expression of miR156 represses nodulation and causes morphological and developmental changes in Lotus japonicus. Molecular Genetics and Genomics 290, 471–484.

Wang X, Yan L, Li T, et al. 2024. The lncRNA1-miR6288b-3p-PpTCP4-PpD2 module regulates peach branch number by affecting brassinosteroid biosynthesis. New Phytologist 243, 1050–1064.

Wang R, Yang X, Guo S, Wang Z, Zhang Z, Fang Z. 2021b. MiR319-targeted OsTCP21 and OsGAmyb regulate tillering and grain yield in rice. Journal of Integrative Plant Biology 63, 1260–1272.

Wang F, Yao T, Yang W, Wu P, Liu Y, Yang B. 2022. Protocol to detect nucleotide-protein interaction in vitro using a non-radioactive competitive electrophoretic mobility shift assay. STAR Protocols 3, 101730.

Wang M, Zang L, Jiao F, Perez-Garcia M-D, Ogé L, Hamama L, Le Gourrierec J, Sakr S, Chen J. 2020b. Sugar Signaling and Post-transcriptional Regulation in Plants: An Overlooked or an Emerging Topic? Frontiers in Plant Science 11, 578096.

Wei S, Gruber MY, Yu B, Gao M-J, Khachatourians GG, Hegedus DD, Parkin IA, Hannoufa A. 2012. Arabidopsis mutant sk156 reveals complex regulation of SPL15 in a miR156-controlled gene network. BMC Plant Biology 12, 169.

Wei H, Luo M, Deng J, et al. 2024. SPL16 and SPL23 mediate photoperiodic control of seasonal growth in Populus trees. New Phytologist 241, 1646–1661.

Weigel D, Ahn JH, Blázquez MA, et al. 2000. Activation Tagging in Arabidopsis. Plant Physiology 122, 1003–1014.

Wu F, Qi J, Meng X, Jin W. 2020. miR319c acts as a positive regulator of tomato against Botrytis cinerea infection by targeting TCP29. Plant Science 300, 110610.

Xie Y, Liu Y, Ma M, Zhou Q, Zhao Y, Zhao B, Wang B, Wei H, Wang H. 2020. Arabidopsis FHY3 and FAR1 integrate light and strigolactone signaling to regulate branching. Nature Communications 11, 1955.

Xie Y, Liu Y, Wang H, Ma X, Wang B, Wu G, Wang H. 2017. Phytochrome-interacting factors directly suppress MIR156 expression to enhance shade-avoidance syndrome in Arabidopsis. Nature Communications 8, 348.

Xing Y, Zhang Q. 2010. Genetic and Molecular Bases of Rice Yield. Annual Review of Plant Biology 61, 421–442.

Xiong Y, McCormack M, Li L, Hall Q, Xiang C, Sheen J. 2013. Glucose–TOR signalling reprograms the transcriptome and activates meristems. Nature 496, 181–186.

Yang Y, Nicolas M, Zhang J, Yu H, Guo D, Yuan R, Zhang T, Yang J, Cubas P, Qin G. 2018. The TIE1 transcriptional repressor controls shoot branching by directly repressing BRANCHED1 in Arabidopsis. (GS Barsh, Ed.). PLOS Genetics 14, e1007296.

Yang L, Xu M, Koo Y, He J, Poethig RS. 2013. Sugar promotes vegetative phase change in Arabidopsis thaliana by repressing the expression of MIR156A and MIR156C. eLife 2, e00260.

Zhan J, Chu Y, Wang Y, Diao Y, Zhao Y, Liu L, Wei X, Meng Y, Li F, Ge X. 2021. The miR164-GhCUC2-GhBRC1 module regulates plant architecture through abscisic acid in cotton. Plant Biotechnology Journal 19, 1839–1851.

Zhang B, Pan X, Cobb GP, Anderson TA. 2006. Plant microRNA: A small regulatory molecule with big impact. Developmental Biology 289, 3–16.

Zhao W, Li Z, Fan J, Hu C, Yang R, Qi X, Chen H, Zhao F, Wang S. 2015. Identification of jasmonic acid-associated microRNAs and characterization of the regulatory roles of the miR319/TCP4 module under root-knot nematode stress in tomato. Journal of Experimental Botany 66, 4653–4667.

Zhao X, Yang J, Wang H, Xu H, Zhou Y, Duan L. 2025. MicroRNAs in Plants Development and Stress Resistance. Plant, Cell & Environment 48, 5909–5929.

Zheng X, Lan J, Yu H, Zhang J, Zhang Y, Qin Y, Su X-D, Qin G. 2022. Arabidopsis transcription factor TCP4 represses chlorophyll biosynthesis to prevent petal greening. Plant Communications 3, 100309.

Zhou M, Li D, Li Z, Hu Q, Yang C, Zhu L, Luo H. 2013. Constitutive Expression of a miR319 Gene Alters Plant Development and Enhances Salt and Drought Tolerance in Transgenic Creeping Bentgrass. Plant Physiology 161, 1375–1391.

