## Supplemental Figure S1 for "Sugar metabolism drives shoot branching by repressing *BRC1* through *miR319*-targeted TCP4 in *Rosa*"

### Supplementary data 1

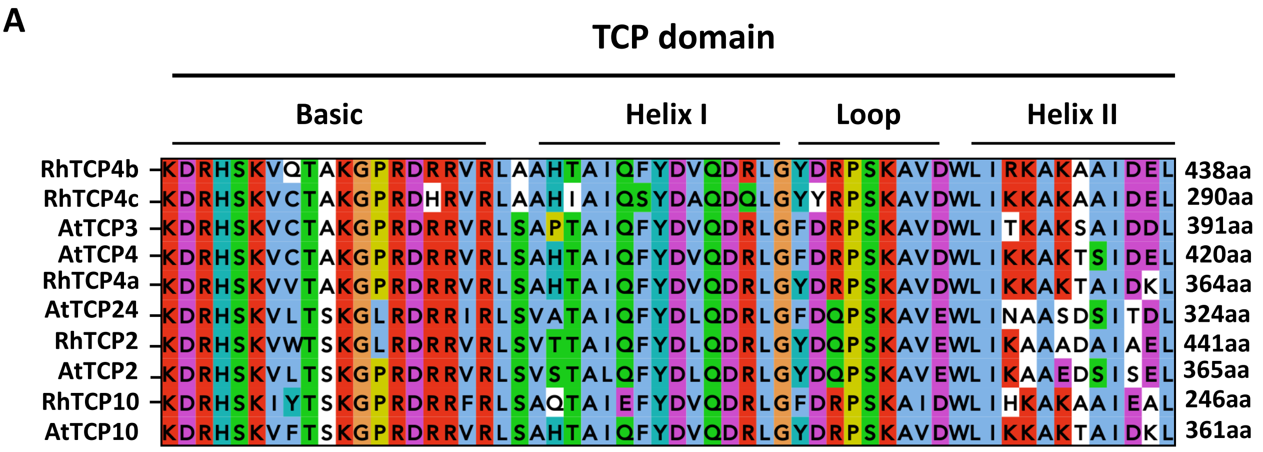


**B**

```
          1                                                                                                                           130
   RhTCP2                                                                     MEVEEIQAQ QPRKFPRIGN GGRGSTIVKS PDEEEDGGRE LKRAGGGDAG TNRLNGWHHS RIIRVSRASG
  RhTCP4a                                                                                                                  MGM KSVGGEIVQV QGGHIVRSTG
  RhTCP4b                                                                                        MGESHNNLHH RATTSSSAAS RLGLGIRTAS SSTTSEIVEA RG-HILRATG
  RhTCP4c MCDARESLNE DKKTDLPSKV FFSFSFLSFF LYSFFFPFLF FSFFLSLFFF FFFLGFFFLS FSPCQHHRFR RTTIEYAGRK MGESHNNLHH RTMTSSSAAS RLDLGIRTAS SSTASEIVEA RG-HILRSTG
  RhTCP10                                    MFVK LERKKRETSM TSHSQESLYL QMSQQRESKQ EEEEEEEEEE ENEAQDQPEV AQAQQDYGAS RVESKQPSMN NERYGRIVKV QGGHVIRSIA
Consensus .......... .......... .......... .......f.. .......... .......... .m..q....q ....e..... .ge....... .....d.gas r...g...a. ..r...iv.. rg.h!.Rs.g

          131                                                                                                                         260
   RhTCP2 GKDRHSKVWT SKGLRDRRVR LSVTTAIQFY DLQDRLGYDQ PSKAVEWLI- --KAAADAIA ELPSLNSSFP DTPKQLSDEK RASSERGFDS VELDVDGDPN YHHQVQNHHQ LCMSKSACSS TSETSKGSGG
  RhTCP4a RKDRHSKVVT AKGPRDRRVR LSAHTAIQFY DVQDRLGYDR PSKAVDWLIK KAKTAIDKLA ELPPWHPTGV AANATQG--- --EVPNGAGT SDMVISAEQS ESSGYNFQLQ RQIGDN---- -TDHNESSFI
  RhTCP4b RKDRHSKVQT AKGPRDRRVR LAAHTAIQFY DVQDRLGYDR PSKAVDWLIR KAKAAIDELD ELPPWNPNSI SVQTTSSPTV PISVAQDTQN ARLHFSMVEA PSSLSANRRG AMVGSSGVAE LVNQNSSNFL
  RhTCP4c RKDRHSKVCT AKGPRDHRVR LAAHIAIQSY DAQDQLGYYR PSKAVDWLIK KAKAAIDELD ELSPWNPNSI SVQIISSPTV PISAAQDTQN ASLHFSMVEA PSSLSANQQG AMVGSNGVAE LVNQNNSNFL
  RhTCP10 RKDRHSKIYT SKGPRDRRFR LSAQTAIEFY DVQDRLGFDR PSKAIDWLIH KAKAAIEALE DSSTTH---- ------QPQH HQGGNAEPET VKTHQPLDEG TMKNESKEPV LDYAEEALNC APNLSEANFM
Consensus rKDRHSK!.T sKGpRDrRvR Lsa.tAI#fY DvQDrLG%dr PSKA!#WLI. kaKAAi#al. #ls..n.... ......sp.. ..s.....#. v.lh...de. ......n... l.....a... ..#.s..nf.

          261                                                                                                                         390
   RhTCP2 LSLSRSEMRV NRDKARERAR ERAAKEKEK- ---DSNNDES SRIAQAISQN ASFTELLTGG IGNNRNNSPT GHQQNNPNGE PILFHKAAPP MDYFGSSLLG LSSTPTRNHH QFSGQIHLGN SLPQTMFSIS
  RhTCP4a PQSLDSDTIA DTMKSFFP-- TSSAASTINF Q--SYPSDII SRTTNLNPNQ DLGLSLHSFQ DHGLIHHTQQ SHQDTTHHND QIQTLFAVPA TAGFDSSA-- ---------- --------YQ RMV-AWSS--
  RhTCP4b PPSMDSDSIA DTIKSFFPMG AATAAETQSF HPASYPPDLL SRTSS--QSQ DLRLSLHSFQ DPILLHHHHQ AHHHHHQTHH ENEQAFLTPN HLGFDSSAAT WTEHHQQQQQ QQQAELNRFQ RMV-TWNNAG
  RhTCP4c QVRMVWRPKL LIFVCLQLPF IEVGLEAEVV G
  RhTCP10 EMGWFQSLMA WNYNAGDGGE GCSFNSSSVT LL
Consensus .........a ....a..... .....e..v. .. ....... .......... .......... .......... .......... .......... .......... .......... ..........

          391                                                                                                                         520
   RhTCP2 GDHNHSHPEM QHFSFVPDYI PVSAAANGGE YNLNFSMSSS GGLAGFNRGT LQSNSSSSPS LLP------- -HLQRFSPMD GSSSVPFFIG AAAAAAAPTM ENHQFPAGLQ LYGDGSRHSD HKGKAKN
  RhTCP4a -DSSHENRG- ------FVFN NMPALPHQ-Q ALMGQQQSSS AAPTFAQRGT LQSSFTPSIR SWSD------ -EIQMASQRT QQHPVHH--S SIFGTRFASD GLPVFCTPTR IHGDDADNSV VSDRPSSTSS
  RhTCP4b GDSNNNNNSS SGGGVEFTFN SQPQTQSQ-S PTLQPFLFGQ NQFFSSQRGP LQSSNTPSVH AWIDHNDLSD HQHQMAPSIH SHHHHHHQQS SISSMGFASG GFSGFHIPAR IQGEE-EHDG ISDKPSSASS
  RhTCP4c
  RhTCP10
Consensus .......... .......... .......... .......... .......... .......... ....          .......... .......... .......... .......... .......... ..........

          521
   RhTCP2
  RhTCP4a PNSNHH
  RhTCP4b -NSRH
  RhTCP4c
  RhTCP10
Consensus  ....
```

TCP domain

*RhmiR319* site

**Fig. S1. Conservation of TCP domain truncation of rose TCP proteins.** Supports Fig. 2.
(A) Alignment of the TCP domains between *Arabidopsis* and rose proteins. The basic regions, helix I, loop, and helix II are indicated. (B) Alignment showing that C-terminal truncation of RhTCP10 and RhTCP4c results in the loss of the *miR319* target site.
