## Supplemental Figure S2 for "Sugar metabolism drives shoot branching by repressing *BRC1* through *miR319*-targeted TCP4 in *Rosa*"

### Supplementary data 2

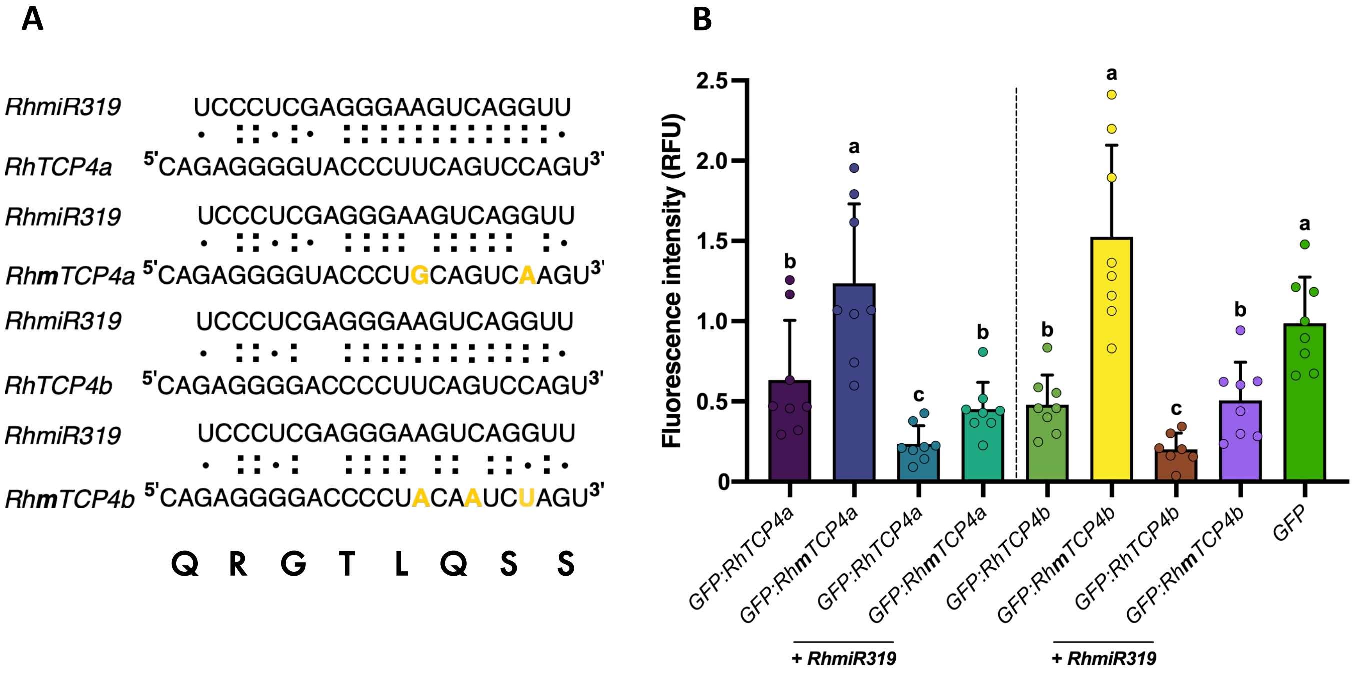


**Fig. S2. *RhmiR319* targets *RhTCP4* genes.** Supports Fig. 2.
(A) Sequence details of *miR319*-binding site in wild-type *RhTCP4a*, *RhTCP4b* and mutated versions (*RhmTCP4a*, *RhmTCP4b*). Mutated nucleotides are shown in orange. (B) *RhTCP4a*, *RhTCP4b*, *RhmTCP4a* and *RhmTCP4b* fused to the GFP reporter gene were co-infiltrated alone or with *35S::RhmiR319* (*RhmiR319*). The GFP fluorescence intensity was measured at the end point 48 hours after infiltration. Data represent the mean ± SEM ($n = 8$). Letters indicate significant differences ($P < 0.05$) based on a non-parametric Kruskal–Wallis test followed by Dunn's post hoc test.
