## Supplemental Figure S3 for "Sugar metabolism drives shoot branching by repressing *BRC1* through *miR319*-targeted TCP4 in *Rosa*"

### Supplementary data 3

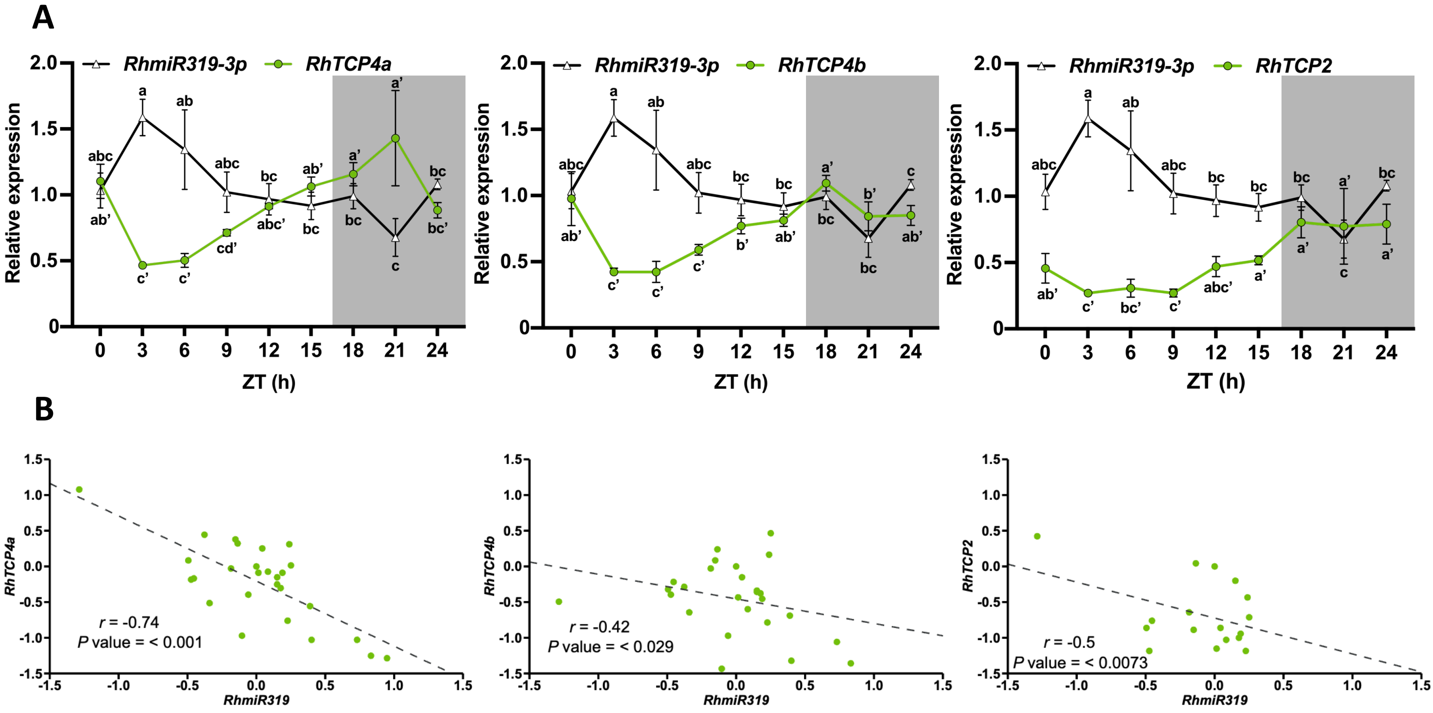


**Fig. S3. *RhmiR319* negatively regulates *RhTCP4* genes.** Supports Fig. 3.

(A) Levels of *RhmiR319*, *RhTCP4a, RhTCP4b*, and *RhTCP2* in axillary buds of intact plants over 24 h. (B) Correlation analyses between *RhmiR319* transcription levels and *RhTCP4a*, *RhTCP4b*, and *RhTCP2* targets in rose axillary buds. Data from the biological replicates shown in (A) were used. The Pearson correlation coefficient (*r*) and probability (*P*) values for each relationship are shown. Data represent the mean ± SD from three biological replicates. Letters indicate significant differences ($P < 0.05$) based on a non-parametric Kruskal–Wallis test followed by Dunn's post hoc test.
