## Supplemental Figure S4 for "Sugar metabolism drives shoot branching by repressing *BRC1* through *miR319*-targeted TCP4 in *Rosa*"

### Supplementary data 4

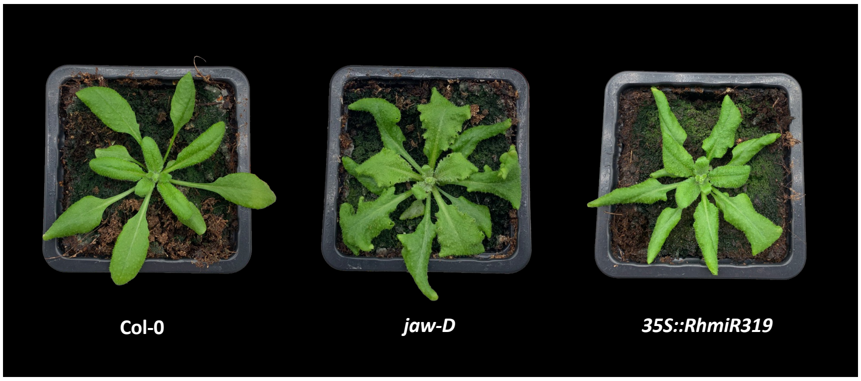


**Fig. S4. Leaf morphology of WT, *jaw-D and 35S::RhmiR319* in *Arabidopsis.*** Supports Fig. 4. Photographs representing leaf morphology of WT, *jaw-D* and *35S::RhmiR319* lines at bolting stage.
