## Supplemental Figure S5 for "Sugar metabolism drives shoot branching by repressing *BRC1* through *miR319*-targeted TCP4 in *Rosa*"

### Supplementary data 5

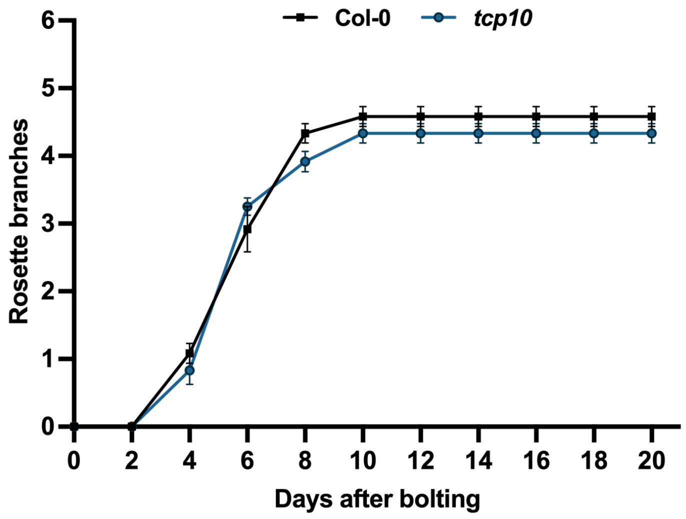


**Fig. S5. Shoot branching in *tcp10* line in *Arabidopsis*.** Supports Fig. 5.
Total number of primary rosette branches in the WT and *tcp10* lines. Data represent the mean ± SEM ($n$ = 12).
