## Supplemental Figure S6 for "Sugar metabolism drives shoot branching by repressing *BRC1* through *miR319*-targeted TCP4 in *Rosa*"

### Supplementary data 6

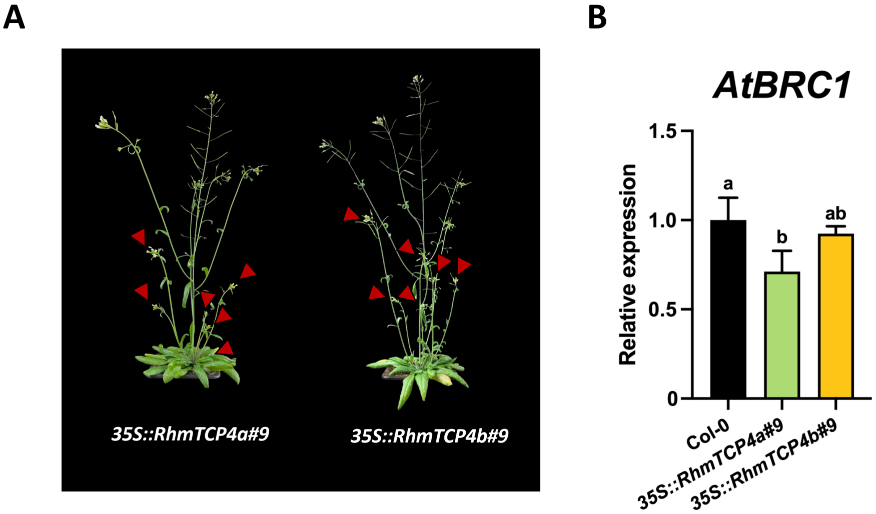


**Fig. S6. Branching phenotype of independent *35S::RhmTCP4a/b* lines in *Arabidopsis*. Supports Fig. 5.**
(A) Photographs representing the branching phenotype of *35S::RhmTCP4a/b* lines at 14 days after bolting. (B) Levels of *AtBRC1* in WT and two other independent *35S::RhmTCP4a/b* lines. Data represent the mean ± SD from three biological replicates. Letters indicate significant differences ($P < 0.05$) based on a non-parametric Kruskal–Wallis test followed by Dunn's post hoc test.
