## Supplemental Figure S7 for "Sugar metabolism drives shoot branching by repressing *BRC1* through *miR319*-targeted TCP4 in *Rosa*"

### Supplementary data 7

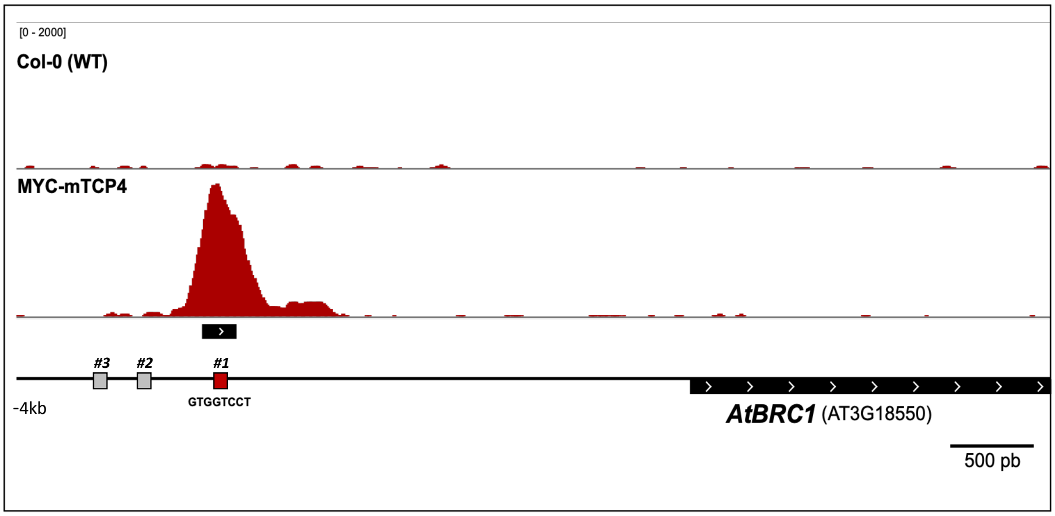


**Fig. S7. TCP4 binds *BRC1* promoter in *Arabidopsis*.** Supports Fig. 6.
ChIP-seq analysis revealed TCP4 enrichment at the *AtBRC1* promoter. Three potential AtTCP4 binding sites in the *AtBRC1* promoter were predicted using JASPAR (red, ChIP-seq peak; grey, predicted sites without ChIP-seq enrichment).
