## Supplemental Figure S8 for "Sugar metabolism drives shoot branching by repressing *BRC1* through *miR319*-targeted TCP4 in *Rosa*"

### Supplementary data 8

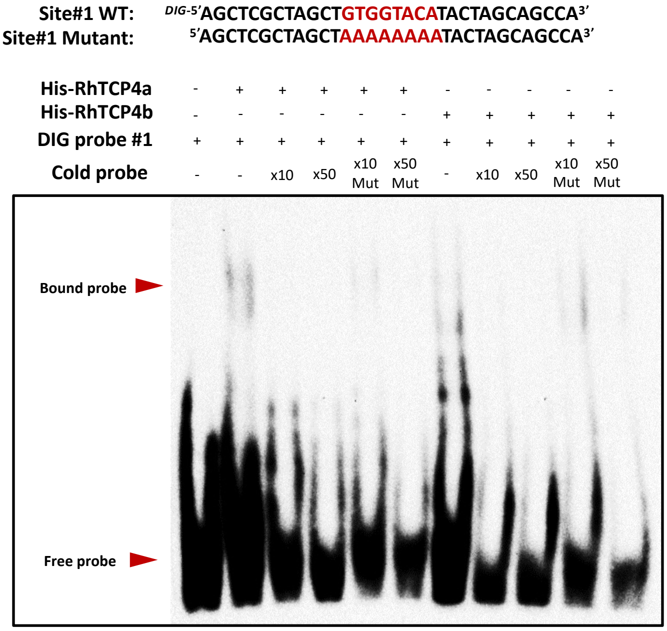


**Fig. S8. EMSA assay using 6xHis-tagged RhTCP4a and RhTCP4b proteins with the site #1 DNA fragment of *RhBRC1* promoter.** Supports Fig. 6.
