## Supplemental Methods S1 for "Sugar metabolism drives shoot branching by repressing *BRC1* through *miR319*-targeted TCP4 in *Rosa*"

**Methods S1. ChIP-seq data analysis**

Raw ChIP-seq data were downloaded from GEO (accession number GSE115589; Dong *et al.*, 2019) and visualized using Integrative Genomics Viewer (v2.16.1). The reads were aligned to the *Arabidopsis thaliana* reference genome (TAIR10) using Bowtie2 (version 2.5.5). Aligned reads were converted to BAM format, sorted, and indexed, and low-quality alignments (mapping quality score < 30) were filtered out using samtools (version 1.23). Coverage profiles were generated using bamCompare from deepTools (version 3.5.6). The ChIP signal was normalized to RPKM (Reads Per Kilobase of transcript per Million mapped reads), calculated over 10-bp bins, and smoothed using a 50-bp sliding window. Background noise was eliminated by mathematically subtracting the control (Input) signal from the target sample signal. Peak calling was performed using homer (version 4.11) with the Input sample serving as the background control. Default stringency parameters were applied, requiring a minimum 4-fold enrichment over the Input (Fold change ≥ 4) and a strict false discovery rate (FDR < 0.001).
