## Supplemental Table S1 for "Sugar metabolism drives shoot branching by repressing *BRC1* through *miR319*-targeted TCP4 in *Rosa*"

**Table S1.** Primers list used in the study.

| RT stem loop | Sequence (5'-3') |
|---|---|
| RhmiR319_RT_Loop | GTCGTATCCAGTGCAGGGTCCGAGGTATTCGCACTGGATACGACGGGAGC |
| RhU6_RT_Loop | GTCGTATCCAGTGCAGGGTCCGAGGTATTCGCACTGGATACGACATTTGG |
| AtmiR319_RT_Loop | GTCGTATCCAGTGCAGGGTCCGAGGTATTCGCACTGGATACGACAGGGAG |
| AtU6_RT_Loop | GTGCAGGGTCCGAGGTTTTGGACCATTTCTCGAT |
| qRhU6 | CGGGCATAAATCGAGAAATGGT |
| qRhmiR319-3p | GCGCGCTTGGACTGAAGGGA |
| qAtU6 | GGAACGATACAGAGAAGATTAGCA |
| qAtmiR319 | GCGGCGGTTGGACTGAAGGGAG |
| qUniversal_Stem loop | GTGCAGGGTCCGAGGT |

| Cloning | Sequence (5'-3') |
|---|---|
| pUPD2_RhproBRC1-PCR1 Fw | GCGCCGTCTCGCTCGGGAGAACTCAAAATGGGGAATGCGA |
| pUPD2_RhproBRC1-PCR1 Rv | GCGCCGTCTCGGGGACGACCGACTCAGAGTC |
| pUPD2_RhproBRC1-PCR2 Fw | GCGCCGTCTCGTCCCTTCGAGTGCAACACTA |
| pUPD2_RhproBRC1-PCR2 Rv | GCGCCGTCTCGCTCACATTTGTGATGTATATAGCTAATATCTGG |
| pUPD2_RhTCP4a Fw | GCGCCGTCTCGCTCGAATGGGCATGAAGAGCGTCGG |
| pUPD2_RhTCP4a Rv | GCGCCGTCTCGCTCAAAGCTCAATGGTGGTTGGAGTTGG |
| pUPD2_RhTCP4b Fw | GCGCCGTCTCGCTCGAATGGGGGAGAGCCACAACAA |
| pUPD2_RhTCP4b Rv | GCGCCGTCTCGCTCAAAGCTCAATGGCGAGAATTGGAGG |
| pENTR_RhmiR319a1 Fw | CACCGTATTAGTGGTATTGAAGCAAGGG |
| pENTR_RhmiR319a1 Rv | TAAGAGGGAATTAGATAAGGGAGC |
| PCR1_RhmTCP4a Fw | ATGGGCATGAAGAGCGTCGG |
| PCR1_RhmTCP4a Rv | GATGGCGGTGTGGGCTGACAACCGGCCGGTGGATCGAACAA |
| PCR2_RhmTCP4a Fw | TTGTTCGATCCACCGGCCGGTTGTCAGCCCACACCGCCATC |
| PCR2_RhmTCP4a Rv | TCAATGGTGGTTGGAGTTGGG |
| PCR1_RhmTCP4b Fw | ATGGGGGAGAGCCACAACAAC |
| PCR1_RhmTCP4b Rv | ATGGCGGTGTGCGCGGCCAGGCGGCCGGTGGCCCTTAGAA |
| PCR2_RhmTCP4b Fw | TTCTAAGGGCCACCGGCCGCCTGGCCGCGCACACCGCCAT |
| PCR2_RhmTCP4b Rv | TCAATGGCGAGAATTGGAGG |

| Genotypage | Sequence (5'-3') |
|---|---|
| LB3_SAIL | TAGCATCTGAATTTCATAACCAATCTCGATACAC |
| LP_SAIL_tcp4 | TTGGGACCAAAAGATTACGTG |
| RP_SAIL_tcp4 | ACTATCATCATCAGCATCCGC |
| LBb1.3_SALK | ATTTTGCCGATTTCGGAAC |
| LP_SALK_tcp10 | TCGAGATCCTTTGGTATCACTG |
| RP_SALK_tcp10 | ATTTCACCACGACTCATTTCG |
