## Supplemental Table S2 for "Sugar metabolism drives shoot branching by repressing *BRC1* through *miR319*-targeted TCP4 in *Rosa*"

**Table S2.** Primers list used for qRT-PCR.

**qRT-PCR**

| Oligo name | ID | Primer | Sequence (5'-3') |
|---|---|---|---|
| **(i) *Rosa*** | | | |
| q*RhUBC* | RC7G0173600 | Forward | GCCACAGATTGCCCATATGTA |
| | | Reverse | TCACAGAGTCCTAGCAGCACA |
| q*RhTCP4a* | RC2G0379100 | Forward | AATGGCTTCACAGAGAACGC |
| | | Reverse | AAACGACGCTGTTGTCTGC |
| q*RhTCP4b* | RC5G0279600 | Forward | AACAATGCGGGTGGTGATAG |
| | | Reverse | AGAACTGGTTTTGGCCGAAC |
| q*RhTCP2* | RC5G0134300 | Forward | TTTGTTCCACAAGGCAGCAC |
| | | Reverse | TTGATGGTGTTTCGGGTTG |
| q*RhASN1* | RC5G0605600 | Forward | CTATTCGAGCCAGCACCCC |
| | | Reverse | TCTCATCAGAGCCCTCACCAG |
| q*RhBRC1* | RC7G0073800 | Forward | TGCATTGTTTAACCCTCTTGCA |
| | | Reverse | GTTCTTTCTCTTGTCTCGCTCTT |
| **(ii) *Arabidopsis thaliana*** | | | |
| q*AtCLATRIN* | AT4G24550 | Forward | AGCATACACTGCGTGCAAAG |
| | | Reverse | TCGCCTGTGTCACATATCTC |
| q*AtTCP2* | AT4G18390 | Forward | ATATCACCGGCAGAATCCA |
| | | Reverse | AGCGAGGAATGACTGATGCT |
| q*AtTCP3* | AT1G53230 | Forward | ATTCGTGCTTGGTTTGATCC |
| | | Reverse | ACCGGAGAATTCACCACTTG |
| q*AtTCP4* | AT3G15030 | Forward | AACGGAGGAGGGTTTCTGTT |
| | | Reverse | TGGAGATGGATTGGTGATGA |
| q*AtTCP10* | AT2G31070 | Forward | TAAGCTTGAACTCGGGGAGA |
| | | Reverse | CACCGTTTGTTGTTGTCGTC |
| q*AtTCP24* | AT1G30210 | Forward | GGACCCTTCAGTCCAATTCA |
| | | Reverse | ATGGTGGTCAAGAGGTGGAG |
| q*AtBRC1* | AT3G18550 | Forward | GATTAACCACCATCGCAGCC |
| | | Reverse | TTTCGCGCCGAAGGAGTAAT |
